# Rough substrates constrain walking speed in ants through modulation of stride frequency and not stride length

**DOI:** 10.1101/731380

**Authors:** G.T. Clifton, D. Holway, N. Gravish

## Abstract

Natural terrain is rarely flat. Substrate irregularities challenge walking animals to maintain stability, yet we lack quantitative assessments of walking performance and limb kinematics on naturally rough ground. We measured how continually rough 3D-printed substrates influence walking performance of Argentine ants by measuring walking speeds of workers from lab colonies and by testing colony-wide substrate preference in field experiments. Tracking limb motion in over 8,000 videos, we used statistical models that associate walking speed with limb kinematic parameters to compare movement over flat versus rough ground. We found that rough substrates reduced preferred and peak walking speeds by up to 42% and that ants actively avoided rough terrain in the field. Observed speed reductions were modulated primarily by shifts in stride frequency and not stride length, a pattern consistent across flat and rough substrates. Modeling revealed that walking speeds on rough substrates were accurately predicted based on flat walking data for over 89% of strides. Those strides that were not well modeled primarily involved limb perturbations, including missteps, active foot repositioning, and slipping. Together these findings relate kinematic mechanisms underlying walking performance on rough terrain to ecologically-relevant measures under field conditions.

## Background

The complex, three-dimensional structures of natural environments disrupt walking at multiple scales. Small-scale surface irregularities (much smaller than body size) reduce foot adhesion, friction, and contact area [1–3], whereas larger-scale environmental structures (those exceeding body size) present navigational challenges [4,5]. Far less is known about how substrate variability at intermediate-scales influences locomotion. Human, animal, and robotic studies on terrain structure at this scale often focus on discrete changes in substrate height, such as crossing a step or gap [6–10]. However, natural substrates present a continuous series of height variations, and animals likely modify their locomotion behavior in distinct ways as they are confronted with these natural substrates. To investigate how intermediate substrate variation influences walking performance, we study one of nature’s most proficient walking animals, the worker ant.

Rough substrates with intermediate-scale structure likely present specific challenges to legged locomotion, but we lack a standardized metric to describe terrain in this context. These substrates are commonly referred to as “rough” and quantified using horizontal and vertical measures of spatial variability [11]. For example, some studies analyze the vertical profile of the ground to determine the average deviation from the mean height or compare the total substrate surface area relative to its horizontal area [12]. Here we define intermediate roughness as substrates with irregularities at the approximate scale of an animal’s body length, and that likely influence stride-to-stride walking patterns. Under these conditions, animals with more than one foot in contact with the ground will likely contend with unequal foot heights, leading to potential pitching and rolling of the body. Roughness may also restrict potential footholds or disrupt walking kinematics.

Ants workers are wingless and walk relatively long distances throughout their adult life [13], thus making them a good subject for understanding legged locomotion over rough terrain. Ants sense and react to properties of the substrate that they traverse, as demonstrated through speed changes on qualitatively rough terrain [14–16] and through optimized foraging pathways that respond to environmental changes [16]. But, the ecological impact of substrate structure likely varies among species. Black garden ants showed no preference for foraging on coarse versus fine sand [15], but seed harvester ants were more likely to drop or transfer seeds when walking on gravel versus sand [17]. These studies suggest that substrate roughness impacts ant foraging performance in some species, motivating analysis that directly ties substrate structure to walking and foraging performance.

Like many insects, ants generally walk using an alternating tripod gait, coordinating the movements of each middle limb with the fore- and hindlimb on the opposing side [18]. To accommodate movement on slopes or when carrying loads, ants can alter limb kinematics by modulating foot placement [19,20], the frequency and timing of stepping [21], and the contact forces produced by each foot [22]. However, similar deviations from stereotypical walking patterns have not been associated with substrate structure in ants. For cockroaches, running over a continuous array of blocks of varying heights reduce speeds by 20% and increase variability in body angular orientations. However, cockroaches do not appear to adjust foot placement of posterior legs to coincide with known good footholds by anterior limbs [23]. To our knowledge, outside of this study on cockroaches, insect limb kinematics has not been analyzed on continuously uneven terrain.

Environmental cues and internal motivational states impact insect behavior [24] representing an important factor for studies on locomotion. Most biomechanical studies account for behavior by inducing a consistent escape response or restricting analysis to a subset of behaviors (i.e. walking in a straight line or at a steady speed). But, quantifying locomotion performance across the natural range of behaviors is critical, especially when deciphering how substrate variability influences walking. Advances in new markerless and automated tracking methods accelerate data acquisition and analysis [25], empowering studies of natural walking that incorporate both environmental and behavioral variation. Using these techniques, we aim to comprehensively quantify normal walking kinematics while embracing behavioral variability.

Here we perform laboratory and field experiments to quantify the impact of uneven terrain on the walking kinematics and preferences of unrestrained Argentine ant workers (*Linepithema humile*). In laboratory experiments we recorded thousands of videos of ants walking on 3D-printed flat and rough substrates with a roughness scale approximately greater than, equal to, and less than worker body length (Fig 1a-c). In outdoor experiments we used the same substrates to test if colonies of ants would demonstrate a preference between rough versus flat substrates. To understand these measures of walking performance, we used a deep-learning approach to track limb kinematics on flat and rough substrates, identifying touchdown locations and timing. Using these measures, we modelled walking speed based on flat ground strides and analyzed how this model predicts speeds on rough ground. Our findings provide the first description of walking kinematics for ants on continuously rough terrain while incorporating variability due to diverse, natural behaviors.

**Fig. 1.**
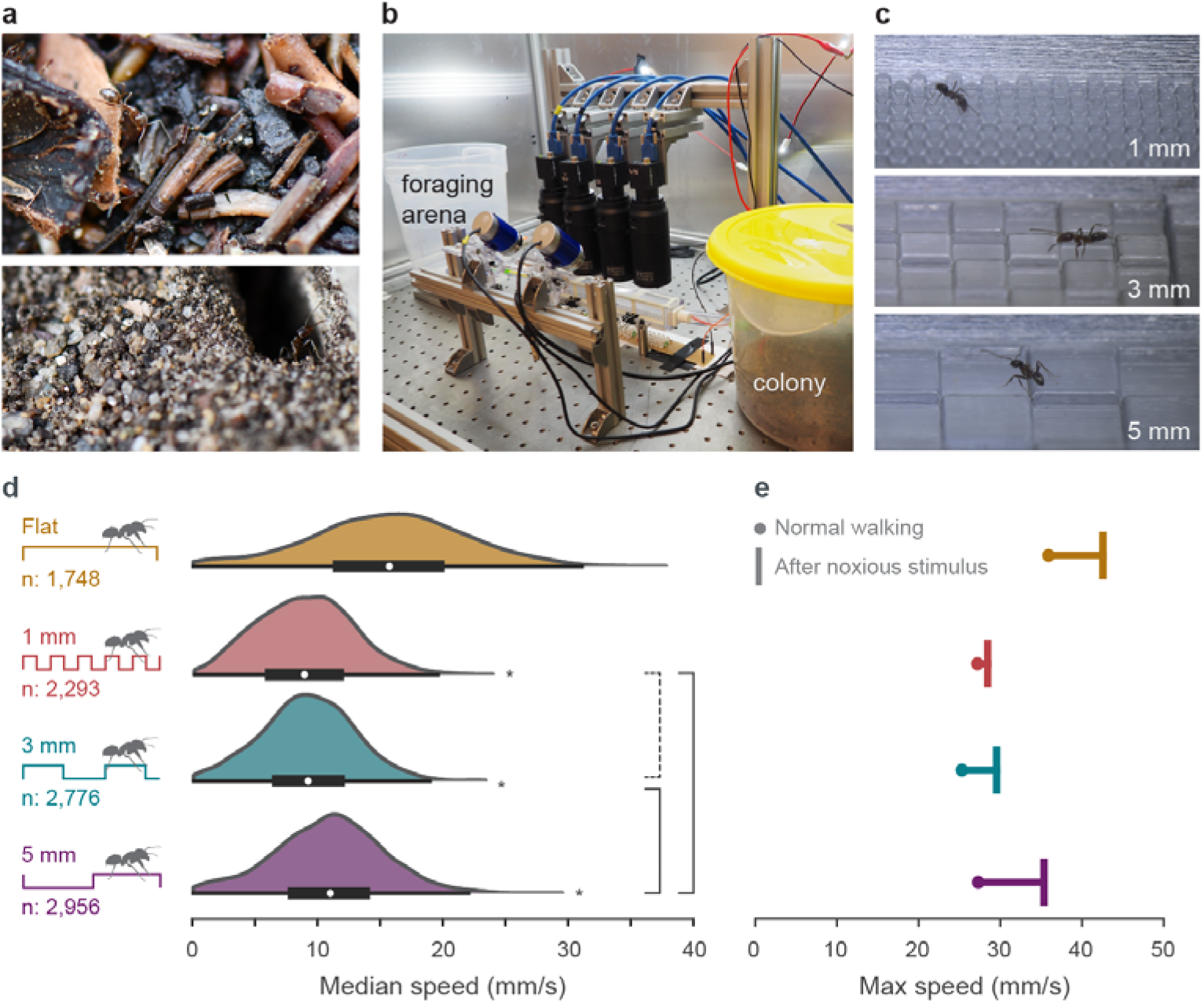
3D-printed rough substrates constrain average and peak walking speeds in ants. (a) Ants encounter uneven natural terrains with roughness features at varying scales. (b) High-speed cameras recorded Argentine ant workers walking on flat and rough substrates. (c) 3D-printed checkerboard patterns with a 1 mm step height and 1, 3, or 5 mm box width. (d) Preferred walking speeds decreased on rough substrates. Speed distributions on all rough substrates significantly differed from flat ground (asterisks, p < 0.0001, see SI for statistical analysis) and from each other (solid brackets, p < 0.0001) except for 1 and 3 mm substrates (dashed bracket, not significant). (e) Maximum walking speeds were compared for unperturbed trials (dot) and those following a noxious airburst (“T”). Peak speed increased the least on the 1 mm substrate and most on the flat and 5 mm substrates.

## Results and Discussion

### Walking speeds in ants from laboratory colonies

We recorded 8266 videos of ants recruiting to food through a tunnel lined with randomly- ordered substrates, including one flat and three “rough” checkerboard patterns (Fig. S1a-d). The checkerboard step patterns had a 1 mm step height and box lengths in three sizes: 1, 3, or 5 mm (Fig. 1c, Fig. S1d). Tracking ant locations across frames (Fig. S2), we find that ants walked more slowly on rough substrates. Instantaneous speed varied within and among steps (Fig. S3), but median speed across the flat substrate was on average faster (15.1 mm/s) than on rough substrates, particularly for the 1 and 3 mm checkerboards (8.7 and 8.9 mm/s respectively) (Fig. 1d, Fig. S4b).

Rough substrates reduce walking speeds in ants with our results clarifying the impact of roughness scale. Ants are known to slow down on rough substrates that qualitatively differ (e.g. gravel vs. sand, medium grass coverage vs. bare ground)[15,17,26–28], but the only previous study that directly associated the scale of surface roughness to walking speed found that ants slow down when walking over particles greater than ⅓ body length [14]. All three rough substrates tested in this study exceed this cutoff (worker body length ~ 3 mm), confirming that roughnesses at scales above ⅓ body length influence speed. However, our results show that preferred speed does not decrease linearly with roughness scale and that walking speed is slowest at the intermediate size roughness on our substrates. With similar average speeds on 1 and 3 mm substrates, there is not a distinct “worst case” roughness scale. It is unclear whether walking speed is a smooth function of ground roughness, or if different roughness scales induce distinct walking behaviors.

Because ants do not walk in straight lines or at steady speeds (Fig. S3), focusing on the median speed across a given trackway masks important details of walking performance. Differences in average walking speeds between substrates could derive from variations in pathway sinuosity, behavioral modulation, or physical speed limitations. To separate these factors, we first isolated sections of relatively straight walking (see Materials and Methods). In straight walking trials we found a substrate-dependent pattern in average speed (Fig. S4c) that aligns with the pattern found when including turning (Fig. S4b), showing that turning is not responsible for speed reductions on rough substrates. Second, we compared the distribution of speeds observed on each substrate to determine whether reductions in average speed resulted from a general shift in speed preference or a transition to burst-and-stop walking. We found a unimodal speed distribution on each substrate (Fig. S4d) suggesting that rough substrates do not cause more frequent accelerations and decelerations, but instead a shift to overall slower speeds. Third, we tested the biomechanical limitations of walking on rough substrates by inducing an escape behavior [29]. A noxious jet of cinnamon-infused air injected into the tunnel every 5 minutes (Fig. S1b) increased peak walking speeds on flat substrates by 6.9 mm/s (19.7%) to speed of 42.0 mm/s (~14 body lengths/s) (Fig. 1e). However, 1 and 3 mm substrates constrained peak speeds to below 30 mm/s, with escape increases of only 1.7 and 4.6 mm/s respectively. Compared to 5 mm substrates with an increase of 8.2 mm/s in peak speed, substrates with an intermediate-scale roughness physically limited peak walking speeds. Together these findings demonstrate that the observed reduction in average walking speed on rough substrates does not result from turning, but instead results from a shift in preferred speed and a constraint on peak speed. Since ant walking speed directly relates to food acquisition rate [27] and the risks associated with foraging duration, such as predation [30] or desiccation [31], substrate roughness likely contributes to colony-level performance.

### Substrate preference in wild ants

To complement our finding that rough substrates constrain walking speeds in laboratory-housed ants, we tested whether ants sense and adjust navigation to avoid less-favorable, rough substrates in the field. We designed an outdoor preference experiment that allowed free-living Argentine ant colonies to recruit to food sources accessed via flat versus rough substrates (Fig. 2a, Fig. S1e, Movie S1). Our experimental design accounted for the influence of pheromone trails, navigational memory, and edge-following behavior [32]. Ants significantly preferred flat substrates over the 1 and 3 mm roughness patterns, but showed no preference on the 5 mm substrate compared to the flat substrate (p<0.001, p<0.01, and p>0.05, two-sample t-test) (Fig. 2b, Fig. S1f). Further, the magnitude of ant avoidance of rough substrates aligned with the observed reductions in walking speed on those substrates (Fig. 1d).

**Fig. 2.**
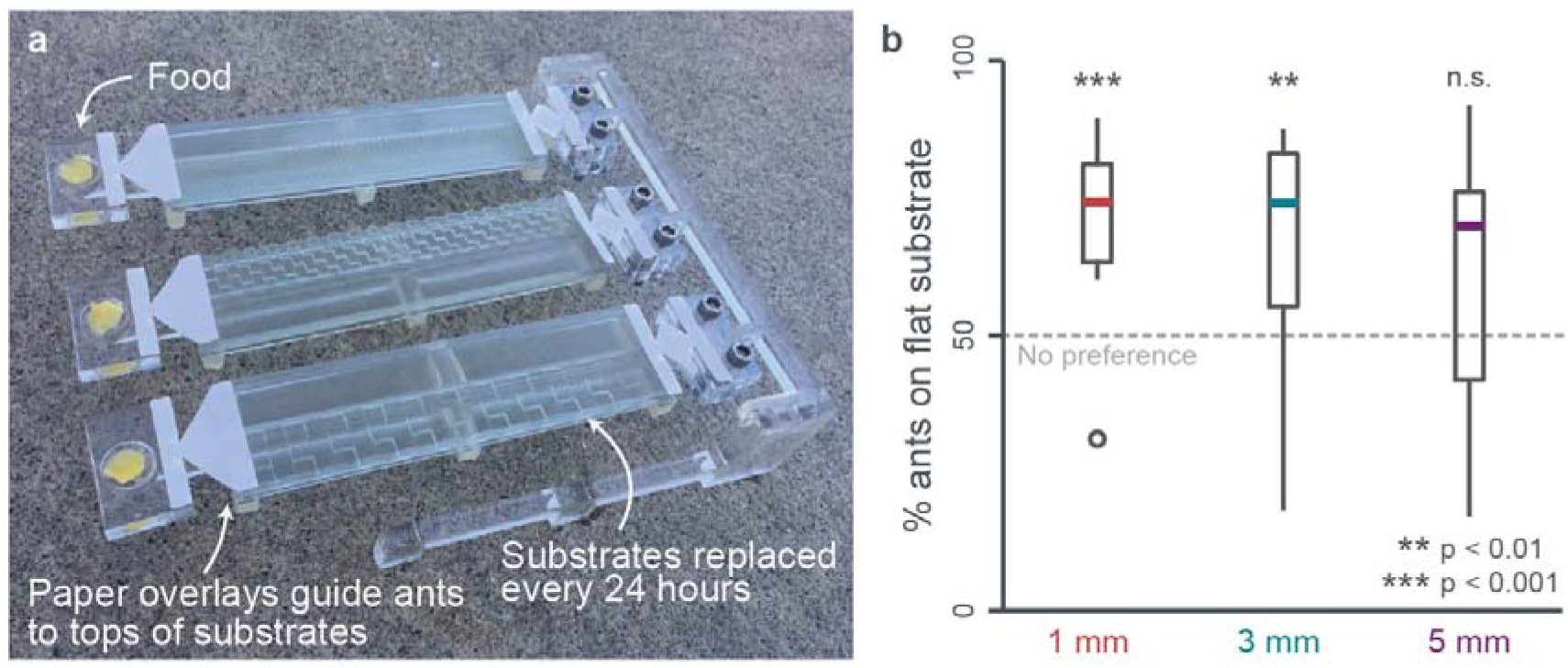
Ants prefer flat ground to 1 and 3 mm, but not 5 mm, checkerboard terrain (see Fig. 1). (a) To test if ants perceive differences in walking performance due to substrate roughness, an outdoor preference experiment tested whether ants would recruit to food sources over flat versus rough substrates. (b) The percentage of ants on flat substrates for each 24 hour trial compared to 50% (dashed line), which represents no aversion to rough substrates. Box-plots correspond to median (center line), upper and lower quartiles (box edges), and data range (whiskers). Significance values for each substrate and for comparisons among substrates are listed in Fig. S1f.

The ability of Argentine ants to perceive relative substrate quality in the field and to adjust navigation accordingly likely contributes to colony-level performance. Previous observations that fire ants converge on trailways that minimize the time needed to walk across a boundary of two rough substrates supports that some ants use speed for path planning [16]. Similarly, path choice experiments in which an ant colony is given the option between two paths of differing length [4,33,34] show a preference for paths that minimize travel time. Here we confirm that when presented with multiple rough substrates Argentine ants select paths that enable faster walking speeds. This finding directly links worker-level walking performance to colony-level foraging behavior, and supports the hypothesis that rough substrates impact colony-level performance. Surprisingly, the selection of “easier” paths is observed even over relatively short walking distances (12 cm) compared to Argentine ant foraging distances in the field (on average >12 m and up to 63 m)[35]. More generally, this sensitivity to substrate structure may provide a critical link between the biomechanical challenges of locomotion on natural substrates and habitat preference in different species.

### Ant limb kinematics

Up to this point we have focused on average measures of walking performance on rough substrates: speed and preference. However, we have not related walking performance to limb kinematics and disruptions from steady walking patterns. Like many insects, ants generally walk using an alternating tripod gait, coordinating the movements of each middle limb with the fore- and hindlimb on the opposing side [18]. To accommodate varying external conditions, such as climbing vertically or carrying a load, ants alter limb kinematics by modulating foot placement [19,20], the frequency and timing of stepping [21], and the contact forces produced by each foot [22]. These shifts in limb motion, coordination, and dynamics offer an opportunity to identify disruptions from normal walking and to test whether these disruptions dictate overall performance on rough substrates.

Traditionally, biomechanical studies of highly-variable limb motion required manual tracking, with the time-intensity of this approach precluding analysis of the large datasets needed to capture the variability of freely-behaving animals. Advances in new markerless and automated tracking methods accelerate data acquisition and analysis, enabling our assessment of walking kinematics in freely recruiting ant workers. To track limb movements in our full data set of videos we used a deep-learning approach. But because walking on rough substrates includes considerable kinematic variability, we performed extensive post-processing to ensure high confidence in the automated tracking (Fig. S6). We analyzed tarsal motion with respect to the substrate to identify touchdown (TD) timing and location (Fig. S7), with sequential TDs defining strides for each limb. Comparing our computational identification of TDs to a subset of manually-tracked videos on each substrate, our approach correctly identified 239/244 TDs (98%), with an average timing offset of less than 1 frame for all substrates. With this approach, we analyzed more than 11,700 walking bouts, corresponding to approximately 2,500,000 video frames.

In legged locomotion (where slipping does not occur) an inter-relationship exists between stride length, stride frequency, and speed. Scaling relations derived and confirmed in vertebrates and several invertebrates [36,37] predict linear relationships in both speed versus stride frequency and speed versus stride length. We confirm a largely linear stride frequency relationship in Argentine ants walking on flat ground, but without an observed plateau at a maximally sustained stride frequency. According to allometric scaling equations [37], this transition should occur in Argentine ants (mass = 0.43 g) at a stride frequency of 25 Hz and speed of 78.6 mm/s, well above the observed data range (40 mm/s). The stride lengths of ants on flat ground primarily range from 1.8 to 2.6 mm, with variation obscuring a clear linear pattern with speed as observed in cockroaches and other ants [18,36]. This discrepancy from other studies does not result from including turns since we restricted our analysis to strides with a heading of less than 15 degrees. However, no observation of a linear stride length versus speed relationship could derive from behavioral variability present in our study that was missing from most prior biomechanical analyses. Alternatively, our findings could represent that ants maintain a relatively constant stride length while walking on flat ground.

To test how stride length versus stride frequency contributes to ant speed, we compared three linear mixed effects models applied to “stereotypical” strides from ants walking on flat ground (Fig. 3a, S8). The “full” model included both stride length (SL) and stride frequency (SF) as factors, with the other two reduced models each lacking one of those variables (“constant SL” and “constant SF”). Comparing model fit between the full and reduced models showed that the constant SL model is 3.5 times more similar to the full model compared to the similarity between the constant SF and full models (chi squared values of 14,644 vs. 53,301 for constant SL and constant SF respectively) (Table S1). While the exclusion of either factor caused a significant difference from the full model (p < 0.001), our large sample size contributes to this finding.

**Fig. 3.**
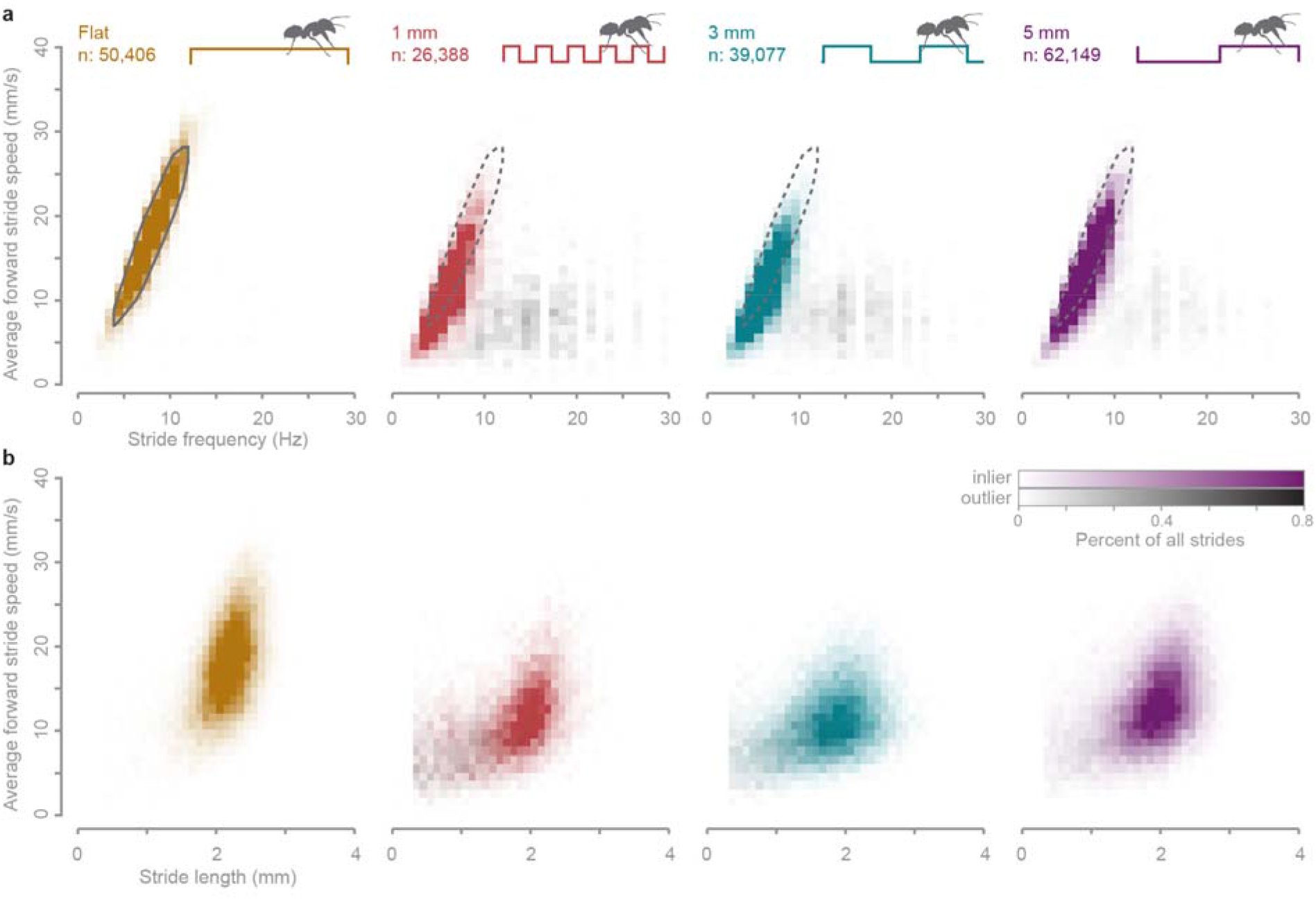
Speed, stride frequency, and stride length relationships for walking on flat and rough substrates. (a) The average instantaneous speed across a stride varies roughly linearly with stride frequency. Stereotypical flat walking strides were identified using a 2D-density cutoff (gray solid and dashed polygons). (b) Stride length remains relatively constant across average stride speeds. For both panels, data plotted in color represents strides well-modeled by the “full” linear mixed-effects model generated from stereotypical flat walking strides (“inlier strides”, see Materials and Methods for details). Gray data represents strides that were poorly-modeled (“outlier strides”).

As a secondary assessment of the full and reduced models, we used each model to predict the stride speed for all strides on all substrates then compared these values to the observed speeds. In addition to evaluating relative model performance, this approach enables comparisons of walking strategies across substrates. The error distributions between the predicted and observed speeds for the full model consisted of two parts, which we isolated to analyze separately (Fig. 4a). A largely normal portion of the distribution defined the “inlier strides,” while a long tail of “outlier strides” contained points that largely deviate from the observed speeds. The percent of strides in each category varied by substrate, with inlier strides representing between 89-99% of all strides (Fig. 4b).

**Fig. 4.**
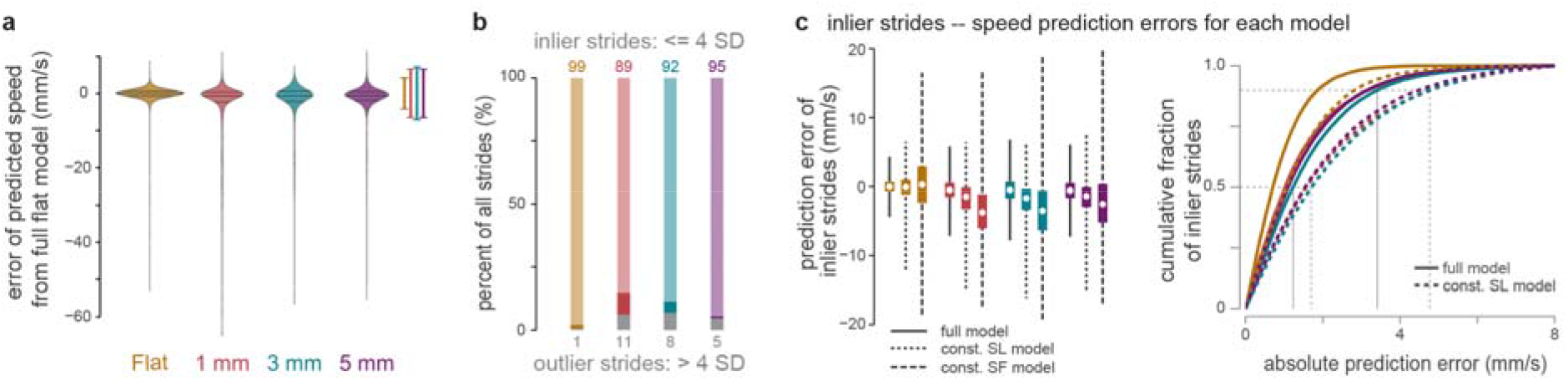
Modeling of walking speed using stride frequency and stride length from stereotypical strides on flat ground. (a) A full model using both stride frequency and length predicted most strides with a small error from observed values. The highest density regions of these distributions were used to determine the cutoff for inlier (brackets) and outlier strides on each substrate. (b) The percentage of strides that were inlier (bars in color) versus outlier (gray bars) for each substrate. The darkest, colorful portion of each represents the range in percentages across all seven lab colonies. (c) The error between predicted and observed inlier speeds for the full model (solid lines), constant stride length model (dotted lines), and constant stride frequency model (dashed lines) on each substrate. Box edges represent quartiles with whiskers representing the full range. A cumulative density plot shows the fraction of strides versus prediction error. Thin, solid (full model) and dashed (constant stride length model) lines represent the prediction error cutoffs for 50% and 90% of strides.

Focusing on the inlier strides, the full model predicted stride speed on average within 0.56 mm/s on all substrates (Fig. 4c). This level of congruity is expected for strides on flat ground, since the model was generated from a large portion of those strides. But surprisingly, the full flat ground walking model also predicted the forward speed of an ant walking on rough substrates within a small margin. Over 50% of the inlier strides on all substrates were predicted within 1.5 mm/s, with 90% predicted within 3.5 mm/s (Fig. 4c). Accounting for the absolute observed speeds for each stride (with most strides between 5 and 20 mm/s on rough substrates), more than half of all inlier strides had a predicted velocity within 10% of the observed velocity. In comparison, the constant SL model predicted stride speed on average within 1.6 mm/s and the constant SF model predictions averaged within 3.9 mm/s. All models performed best for strides on a flat substrate and worst on 1 mm substrates. The constant SF model did not predict stride speed as well as the constant SL model, as demonstrated by a larger median and broader range in prediction error (Fig. 4c). The constant SL model predicted over 50% of all inlier strides within 1.8 mm/s and 90% of strides within 4.8 mm/s (Fig. 4c), compared to 4.3 mm/s and 9.0 mm/s respectively for the constant SF model (Fig. S9).

The accuracy of simple models in predicting stride speed based on stride frequency and/or stride length reveals two main findings of this study. First, we found that stride frequency predominantly dictates stride speed in ants, with stride length remaining relatively constant. Second, we found that most strides on rough substrates accurately resemble walking on flat ground. Previous observations in ants and other insects show that stride length increases linearly with speed, though at a slower rate than stride frequency [18,21,36]. Here we confirmed and extended this pattern, showing that stride frequency alone is sufficient in predicting stride speed despite a low correlation between stride frequency and length (Table S1). Therefore, ants walking on rough substrates shifted to slower stride frequencies, accounting for the observed slower speeds.

Surprisingly, the dependence of speed on stride frequency and not stride length was retained for walking on uneven ground in ants. When confronted with uneven terrain, humans generally reduce step lengths [38,39], particularly in older adults or in individuals with compromised gaits who must compensate for reduced balance [40,41]. Although ants slowed down on rough substrates, these speed reductions were primarily modulated by changes in stride frequency. It is unclear exactly why ants decrease stride frequency on rough ground though this could derive from both biomechanical or behavioral factors. Biomechanically, ants that attempt to move limbs at higher frequencies could experience detrimental foot-ground interactions that disrupt walking, such as foot slipping or missteps, which result in lower stride frequencies. Behaviorally, ants may decrease stride frequency on rough terrain to provide more time for sensory feedback or to slow limb motions in an attempt to proactively avert disruptive foot-ground interactions. Such proactive modulations of frequency may be a plastic response based on previous experience, which could be observed in future studies of learning trials or transitions from flat to rough ground.

### Characterization of highly divergent strides

The full model generated from stereotypical strides on flat ground accurately predicted the average speed for most strides on all substrates, but a portion of strides strongly deviated from the model predictions. These outlier strides only represented 1% of the strides on flat ground, but ranged up to 11% on 1 mm substrates (Fig. 4b). Accounting for six limbs, an 11% outlier rate corresponds to an average of at least one outlier step every two strides. These strides generally have a slow average speed and short duration (high frequency) (Fig. 3, gray), but what walking behaviors cause outlier strides? We compared foot touchdown position for inlier and outlier strides on each substrate, finding highly consistent foot placement on flat ground during inlier strides (Fig. 5a). Surprisingly, the preferred inlier locations were tightly conserved across flat and rough substrates despite increased variability on rough terrain (Fig. 5a,b). However, foot placement in outlier strides on all substrates shifted posteriorly and medially (Fig. 5b), coinciding with the swing trajectory of each foot. The average placement of the feet during outlier strides diverged from the inlier preferred positions by 0.34-0.45 mm (Fig. 5c). Using a logistic regression, we predicted the likelihood of outlier strides based on foot displacement (Fig. 5d). To be identified as an outlier, strides required a larger displacement on flat ground compared to on rough substrates. The displacement pattern among rough substrates mirrored the patterns in both speed reduction and outlier stride frequency, with the most extreme condition occurring for the 1 mm checkerboards, while the 5 mm checkerboards approached flat ground values. These findings confirm that most strides on rough substrates resemble walking on flat ground, with those strides that diverge from this walking pattern associated with touching the foot down away from the normally preferred positions.

**Fig. 5.**
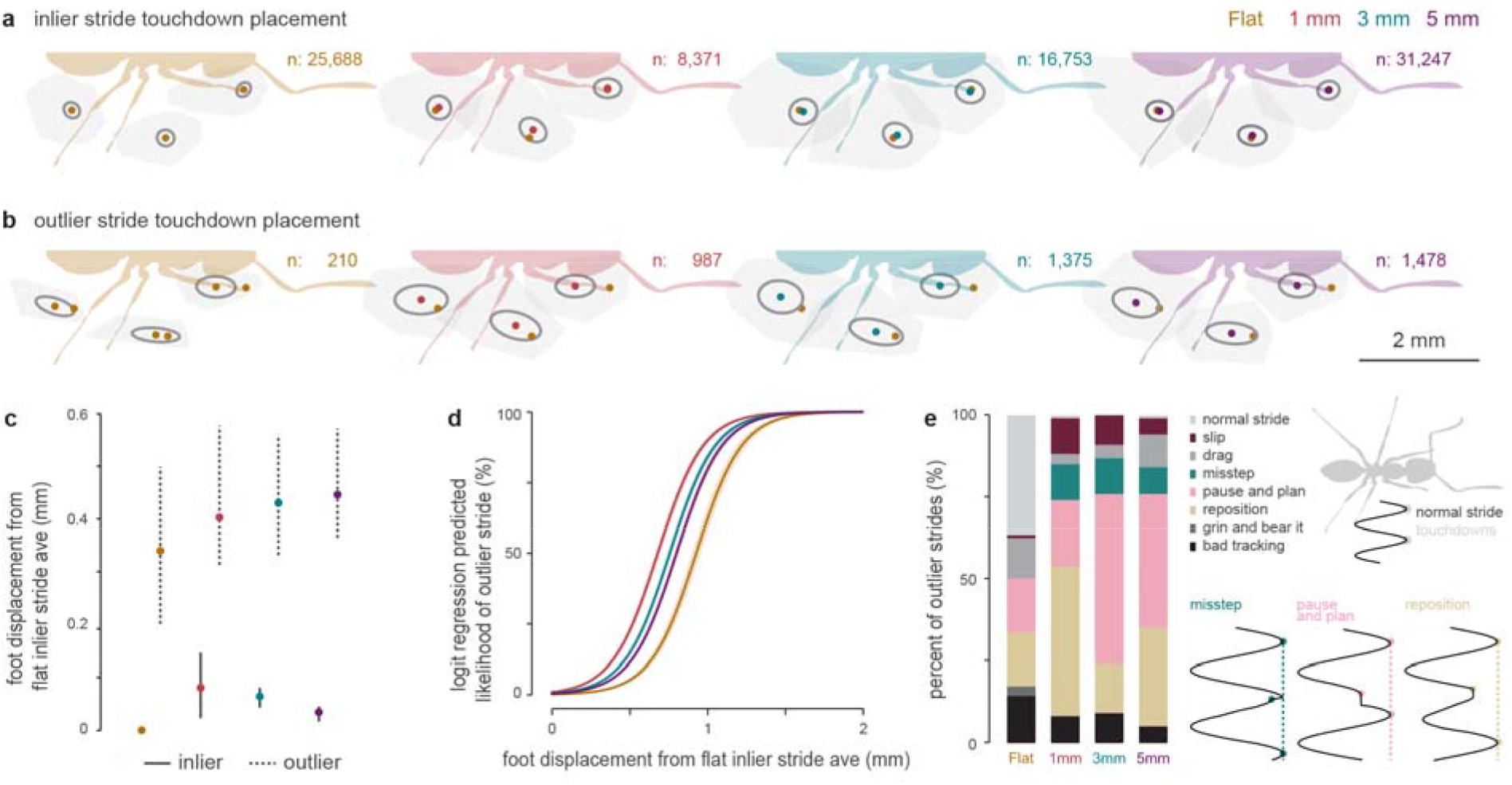
Touchdown foot placement and manual characterization of outlier strides. (a) Average placement of the foot at touchdown for inlier strides on each substrate. Circles represent one standard deviation of the 2D distribution. Yellow data points represent the average location on flat ground. (b) Average placement of the foot at touchdown for outlier strides strides on each substrate. Circles represent one standard deviation of the 2D distribution. Yellow data points represent the average location of inlier strides on flat ground. (c) Total displacement of average foot position relative to average location of inlier strides on flat ground. Whiskers represent the range across limbs, circles represent averages. (d) Likelihood of identification as an outlier stride versus displacement for each substrate. Curves were generated using a logistic regression. (e) Manual characterization of outlier strides on flat and rough substrates. Diagrams represent the anterior-posterior trajectory of the tarsus with respect to the ant (x-axis) across time (y-axis). Dots represent touchdowns.

The “divergent” strides identified through our modeling could result from multiple kinematic behaviors such as slipping [42], premature foot touchdown [43], and active repositioning of the limb during stance for a more favorable foothold [44,45]. To identify the behaviors underlying divergent strides, we manually characterized 100 randomly-selected outlier strides for each substrate (Fig. S10). This characterization represents a first qualitative assessment of outlier strides, motivating future quantitative analyses using gait phasing [9]. We find that on flat ground, with only 210 identified outlier strides, more than half corresponded to normal walking or automated tracking errors (Fig. 5e). Since the tracking error strides represent less than 0.06% of all flat strides, we do not believe that they invalidate our methods or general findings. The remaining flat outlier strides mostly consisted of dragging the foot forwards during stance, repositioning of the foot after a collision, or a brief premature touchdown of the foot as the ant slows down and appears to choose its next step (“pause and plan”). On rough substrates, outlier strides also included missteps, when the foot touches down after reaching and missing its targeted foot placement. Notably, walking on the 3 and 5 mm substrates induced a high percentage of pause and plan strides. These strides mostly coincided with the ant reaching the edge of a checkerboard box and tapping the foot (often of the forelimb) along the corner before placing it forward. Insects possess proprioceptive sense organs within each limb, including both muscle strain sensors and campaniform sensilla in the cuticle, which have been shown to coordinate limb movements, control posture, and detect foot attachment to the ground [46,47]. Most of the studies on insect proprioception focus on insects in controlled conditions or walking on flat ground. The pause and plan strides identified here represent a potential mechanism for ants to use proprioception while walking on uneven terrain.

### Conclusions

Our findings demonstrate that ants are able to mostly conserve flat ground walking patterns and maintain preferred foot placement while on naturally rough substrates. Even though ants reduced walking speeds substantially on rough substrates, over 89% of their strides were accurately modeled based on flat walking data with speed primarily governed by stride frequency and not length. To diminish the influence of the substrate irregularities, ants may use specialized adhesive tarsal pads that secure foot attachment [22] and benefit from polypedalism, which improves passive recovery from impulsive disruptions [8,9,48]. With neuromechanical and robotic control research often focusing on accurate foot placement [49] or navigational planning [50], our findings highlight the potential for developing decentralized mechanisms for navigating uneven terrain. Further, we show that ants in the field seek out substrates that enable faster walking speeds. By extending the large-scale, automated approaches established in this study to multiple species and field conditions, it should be possible to uncover how walking strategies in ants contribute to foraging patterns, habitat selection, assemblage structure, and evolutionary diversification.

## Materials and methods

### Experimental Design: General set-up, ant collection, and recording

To determine how the Argentine ant walks on rough ground, we recorded workers from lab colonies walking over 3D-printed substrates. Between March and May of 2018, material for lab colonies was collected from nine locations on the campus of the University of California, San Diego, with each location separated by at least 500 m (Fig. S1a). After finding a large aggregation of ants (>1000 individuals, including some workers carrying brood), we excavated ~300-600 workers, along with brood and queens, and place them in a lidded plastic container. The data from one colony was removed from analysis since many ants escaped during the recording session.

Lab ant colonies were housed in a container within a custom recording enclosure on a 12:12 hour light:dark cycle (for details, see SI Appendix A). To reach food, ants walked through a 3D-printed tunnel (Connex3 Objet 350, VeroClear material, stratasys Inc., USA) (Fig. 1b) with floor openings for four 3D-printed substrates randomly positioned within the tunnel (Fig. 1c). The substrates measured 16×30 mm and consisted of a flat surface and three checkerboard patterns of a 1 mm step height and a square edge of 1, 3, or 5 mm (SI Appendix A)

The ceiling and side of the tunnel incorporated microscope coverslips (24×50×0.25 mm, AmScope, Ltd., USA) to enable filming. Four machine vision cameras (Blackfly S 13Y3M, PointGrey, Inc., Canada) and lenses (20-100mm, 13VM20100AS, Tamron Co., Ltd., Japan) recorded from above the tunnel. Two webcams (YoLuke A860-Blue, Jide Technology Co., Ltd., China) filmed from an oblique overhead angle to identify when ants approached a substrate.

The tunnel was backlit using a custom array of infrared LEDs (940 nm). During two four-hour recording sessions for each colony, the machine vision cameras recorded ants for 3 seconds at 240 fps (720 frames total, 1000 x 550 pixels). After recording, each machine vision camera paused for 40 seconds to reduce the probability of re-recording the same individual. In total, 8266 videos were recorded across eight colonies.

### Experimental Design: Noxious stimulus procedure

To elicit maximal walking performance, one colony was exposed to a repetitive noxious stimulus. Based on a study that found an aversion to the scent of cinnamon in ants [51], we built a custom system to inject cinnamon-infused air into the tunnel. Pressurized air (15 PSI) was infused with cassia cinnamon oil (Healing Solutions, LLC., USA). The opening of an arduino-controlled solenoid valve allowed the cinnamon-air to flow through a manifold to four tubes inserted into the wall of the tunnel above each substrate. Each noxious stimulus consisted of two, 0.25 second bursts separated by 1.00 seconds. After unperturbed walking was recorded for an hour during each session, cinnamon bursts were triggered every 5 minutes throughout the remainder of the recording session.

### Tracking: Automated full-body tracking

Tracking ant whole-body motion was achieved by identifying all ants in each frame and associating individuals across frames. To identify ants in each frame, we (1) normalized the video using background division, (2) used image processing to isolate the body of each ant, and (3) fit contours to each ant body to estimate the location and orientation (Fig. S2). For details on each step, see SI Appendix B. Ants identified in each frame were associated across frames using a Kalman filter [52].

### Analysis: Average ant speed post-processing and statistical analysis

In Python, the x- and y-coordinates of the kalman-associated ant “trackways” were filtered using a low-pass butterworth filter (Scipy, n=2, □_n_ = 0.2) and differentiated to calculate the instantaneous velocity. Any data points where the center of the ant was within 60 pixels of the edge of the frame were removed. To estimate the average walking speed during a trackway required accounting for behavioral variation, such as antennal cleaning or ant interactions. To identify and remove these stationary behaviors, we calculated the net distance traveled over 90 frames (0.375 s) centered around each time point and removed any datapoint when the ant had moved less than 50 pixels (1.56 mm) during those 90 frames (Fig. S3). This cut-off was manually validated in 20 trials.

Median walking speeds were calculated using these processed speed profiles (Fig. S3). The distribution of median speeds was compared across substrates using linear mixed-effects modeling in R (Fig. 1d, details in SI Appendix C). Briefly, a chi-squared likelihood ratio test determined whether substrate type significantly impacted walking speed and then estimated marginal means of the model were used to calculate the significance of comparisons among substrate types.

Slower walking speeds could result from turning more frequently or more acutely on one substrate. To account for this potential compounding factor, speed was also analyzed for straight walking strides only (with a heading < 15°, see below on “Touchdowns and strides”). The net distance traveled during each straight stride was divided by the stride duration. We applied the above linear mixed-effect model approach to the straight stride dataset (Fig. S4).

### Analysis: Noxious Stimulus

Comparing median speeds for each trackway did not fully encapsulate walking performance. To further clarify how rough substrates impact walking, we calculated the distribution of walking speeds used by ants on each substrate. We applied this analysis to the eighth collected colony, including ants walking before and after a noxious burst of cinnamon-infused air. We accounted for within-step accelerations by calculating a windowed average speed (Fig. S3). Instead of using a time-based histogram of the instantaneous speeds on each substrate, which over-represents slow speeds, we compared the distance traveled at each speed. We restricted our analysis to trials with at least 50 frames (0.21 s) of processed data.

The distribution of distances traveled vs. speed was compared for trackways during the first hour of recording on each day (no noxious stimulus), for trackways recorded less than 2 minutes after a cinnamon airburst, and for trackways recorded between 2 and 5 minutes after a cinnamon burst (bursts were every 5 minutes) (Fig. S5). For flat / 1 mm / 3 mm / 5 mm substrates, the total traveled distances compared were 295 / 485 / 378 / 128 cm. The median speed was identified by finding the speed cut-off with 50% of the total distance traveled at faster speeds. Peak speeds were identified using speed cut-offs of 5% and 2%.

### Experimental Design: Outdoor Preference Experiments

To test whether ants demonstrated an awareness of and preference for substrate roughness, we conducted choice experiments in two locations near the UCSD campus in La Jolla, CA (Fig. S1a). Long rectangular sections (120 x 15 mm) of each substrate type (flat, 1, 3, and 5 mm) were 3D-printed and attached to a custom cantilever-support structure that held 3 pairs of rectangles (Fig. S1e, see SI Appendix D). Each pair of rectangles compared a flat vs. rough substrate, therefore, each set-up tested 1, 3, and 5 mm substrates simultaneously. The end of each pair of substrates contained ant food based on the Bhatkar-Whitcomb diet, combining hard-boiled egg yolk, sugar, and water [53]. To discourage ants from walking on the side of the cantilever support structure or randomly favoring one substrate, a diamond-shaped piece of paper was attached on top of the plank leaded to each pair of substrates.

We recorded ant substrate preference using the cantilever set-ups for 3 weeks during October and November of 2018. A time-lapse camera (Re 16MP camera, HTC, Taiwan) positioned above each set-up recorded images every 3 minutes. After 24 hours of recording, each set of substrates was replaced (in randomized order) before being soaked in a mixture of windex and water for the next 24 hours. This mixture was used to remove all pheromone trails from the substrates before the next recording session. The timelapse videos of the outdoor preference set-ups (Fig. S1e) were automatically analyzed to compare the number of ants on flat versus rough substrates. The total percentage of ant pixels on smooth vs. rough substrates was calculated and compiled for each day of recording (for details, see SI Appendix E).

Ants did not discover the outdoor set-up three times, likely due to the onset of cooler conditions in November. Of set-ups that were discovered, the food sources were discovered 93% of the time, with ants ignoring the substrate bridges once on 3mm bridges and twice on 1 mm bridges. Across both set-up locations, ant preference was tested 13, 14, and 15 times for 1, 3, and 5 mm substrates respectively. For each substrate comparison, we performed a two-tailed T-test to determine whether the percent of ants choosing to walk on a flat vs. rough substrate differed from no-preference, 50% (scipy.stats.ttest_1samp) (Fig. S1f). To test if ant preference differed between each combination of rough substrates, we calculated a two-tailed Welch’s T-test for two independent samples with unequal variances and sample sizes (scipy.stats.ttest_ind).

### Tracking: “LEAP” deep-learning tracking

Manually tracking the limbs in 8,000+ videos is not feasible. Instead, we used a recent deep-learning approach, implemented in MATLAB and Python [25]. The LEAP deep-learning workflow tracks a set of user-defined points on square videos centered on a subject. The user determines a skeleton of connected points to track then iteratively hand-labels frames and predicts tracked point locations on new frames to eventually build a robust training set of labeled frames for the final prediction model. Our training set of 679 manually tracked frames generated a model to predict 10 body landmarks, including the location of each tarsus, in novel frames. This approach worked extremely well for ants walking on flat substrates, but decreased in accuracy for the variability associated with rough substrates. In order to implement this process with our highly-variable data, we (1) produced trustworthy input ant-centered videos, (2) hand-digitized an extensive training set of 679 frames, and (3) implemented post-tracking confidence checks to improve or remove untrustworthy data. Details of these methods can be found in SI Appendix F.

By visually inspecting the raw LEAP and post-processed tracking for 30 recorded videos, we confirmed that our approaches were conservative in removing any potentially inaccurate tracking. In total, 11,700 trackways--corresponding to approximately 2,500,000 video frames--were passed through the LEAP tracking and custom post-processing workflow.

### Tracking: Touchdowns and strides

The deep-learning-based LEAP tracking resulted in time-varying locations of the thorax, neck, and tarsi. These data were used to identify when each foot touched-down (TD) onto the substrate and the strides between these TDs. Because the movement of the body during stance often occluded fore and mid-limb toe-off, we were unable to reliably determine stance and swing timing.

The timing and location of TDs were determined using tarsal speed, but kinematic variability and discontinuous tracking (particularly on rough substrates) required additional measures to ensure accurate TD detection. For details see SI Appendix G. To test the accuracy of this approach, we compared our computationally-identified TDs to visually-identified touchdowns in 18 videos, corresponding to over 57 TDs per substrate type for a total of 244 TDs. 239 of these TDs (98%) were correctly identified within 10 frames, with an average offset of 1 frame (4.2 ms). Of the five TDs identified computationally but not manually, three were confirmed having been missed during the initial manual tracking. The remaining two were false identifications by our post-processing approach.

Strides were defined as occurring between two trusted TDs with two conditions to ensure no intermediate TDs were missed (see SI Appendix G). We calculated four variables for each trusted stride. (1) *Stride frequency* was calculated from the inverse of stride duration (the number of frames between TDs). (2) *Stride length* was defined as the distance between the location of the foot at the beginning and ending TDs. (3) The *average stride speed* was the mean of the instantaneous velocity’s forward component (aligned with the ant’s anterior-posterior axis). (4) The *travel direction* of the ant was the net angular displacement of the thorax of the ant during the stride relative to the ant’s facing at the beginning TD. All strides with a travel direction < 15° were classified as straight walking.

### Analysis: Average stride speed modeling

To test whether ants use stride frequency or stride length to modulate walking speed, we compared three linear mixed-effects models (for details see SI Appendix H, Fig. S8). The source data were drawn from straight strides on flat ground, while removing unusual strides. The full model included both stride frequency and stride length as fixed effects to explain average stride speed. Two reduced models either held frequency or length constant. Each model was used to predict the average stride speed for all strides on all substrates, then we analyzed the errors between predicted and measured speeds. The most dense points of the full model error distribution were used to define a normal distribution, separating inlier and outlier strides with a cutoff of 4 standard deviations (see SI Appendix H). The inlier stride errors were compared using boxplots and an empirical cumulative distribution function.

Foot placement at touchdown was compared for inlier and outlier strides on each substrate. To find the average placement and its variance, we used a principal components analysis (sklearn.decomposition.PCA). The standard deviation of the first two principal components defined the ellipses in Fig. 4D,E. For the first TD of every stride, we calculated the net displacement from the preferred foot placement of inlier strides on flat ground. A logistic regression (see SI Appendix H) was used to predict the probability that a stride was an outlier based on its displacement.

To identify the behaviors associated with outlier strides on each substrate, we manually observed and classified a random subset of these strides. We randomly selected 100 outlier strides from each substrate type and which of eight stride classifications. Two categories involved false identification (normal strides or bad tracking), two represented disruptions during the stance phase (slips and drags), and four represented disruptions during the swing phase (misstep, pause and plan, reposition, grin and bear it) (Fig. S10).

### Data and Code Availability

The datasets generated during and analyzed during the current study are not publicly available due to storage and format restrictions, but are available from the corresponding author upon reasonable request. Code used for analysis is available at https://github.com/gtclifton

## Acknowledgments

We thank Prof. Michael Tolley for 3D printer access and Ben Shih and Chris Cassidy for printing help. We thank Axel Qin for help in experiment prototyping.

## Authors’ contributions

All authors contributed to the conceptualization and methodology design for this study. GC performed data collection, analysis, and drafted the manuscript. All authors contributed to critically revising the manuscript. All authors gave final approval for publication and agree to be held accountable for the work performed therein.

