## Supplementary text for "Rough substrates constrain walking speed in ants through modulation of stride frequency and not stride length"

**Affiliations:**
[1]Department of Mechanical and Aerospace Engineering
University of California, San Diego.
[2]Division of Biological Science
Section of Ecology, Behavior and Evolution
University of California, San Diego.

**This PDF file includes:**

Supplementary text
Figures S1 to S10
Table S1
Legends for Movies S1 to S2
SI References

**Other supplementary materials for this manuscript include the following:**

Movies S1 to S2

### 1. Supplementary Methods

Appendix A: Set-up and recording details

In the lab, ant colonies were kept in a covered plastic container within a custom enclosure made from aluminum framing (80/20 Inc., USA), aluminum sheeting, and blackout curtains. LED strip lights illuminated the experimental enclosure on a 12:12 hour light:dark cycle. A separate plastic bucket contained food (sugar, water, and polymer crystals to delay evaporation).

Ant colonies were collected between 8:00 and 10:00 AM then allowed to acclimate for eight hours in the lab before opening the pathway through the tunnel to the foraging arena (Fig. S1C). After opening the foraging arena, the set-up was not disturbed for the remainder of the experiment. Overnight, the ants explored the foraging arena and developed a recruitment trail through the tunnel. Each colony was then filmed during two recording sessions, from 8:00 AM - 12:00 PM on subsequent days. The ordering of the substrates remained the same for both recording sessions. After the second recording session, the colony was released back to its collection location and the tunnel and substrates were cleaned in warm, soapy water. The equipment dried overnight, dissipating any collected pheromone trails before the first recording session for the next collected colony.

The rough substrates included three checkerboard sizes of a 1 mm step height: 1, 3, and 5 mm box edges corresponded to less than, approximately equal to, and larger than the worker body length (~3 mm). Due to the resolution of the 3D-printer, the edges on the top of each checkerboard box were not exact right angles (Fig. S1D). Measuring the flat portion of two checkerboard boxes for each rough substrate confirmed that the top box edges included a 0.2 mm radius of curvature (ImageJ). Due to this, the flat portions of the top edges were 0.6, 2.6, and 4.6 mm in the 1, 3, and 5 mm substrates respectively. While the relative proportion of roundedness varied with checkerboard size, we believe that the consistency in the absolute measurements provides a uniform challenge across substrates and that the curved edges better reflect natural structures.

Appendix B: Processing of videos to automatically track ant location

To identify ants in each frame we:

(1) We calculated a background image for each video by taking the median of every 100th frame (7 frames total) for each pixel. The background was removed from the video by dividing each frame by the background image.

(2) Each frame was thresholded to keep any pixels with a normalized value greater than 1.2 or less than 0.8. We applied a distance transform to the resulting binary image removing any pixel within 2 pixels of the background then morphologically closed the image using a rectangular structuring element of 5 pixels for 7 iterations. These transformations (a) ensured that the body was identified in one piece (no separation at the petiole or thresholded holes in the abdomen, which is somewhat transparent after an ant drinks water), and (b) removed all limbs and antennae so that ants near each other were correctly separated.

(3) We fit external contours to the thresholded image (OpenCV), removing all contours that were (a) <200 pixels in area, (b) > 1800 pixels in area, or (c) with a centroid within 60 pixels of the image edge. An initial angle of each ant in the image was output from the contour fitting.

To associate identified ants across frames, we used a Kalman filter with inputs of the value and change in the ant contour centroid location, angle, and area. The initial covariance was 0.2 with a state covariance of 1, a max covariance of 10, and max velocity of 100. The accuracy of these parameters was visually confirmed in 20 trials.

### Appendix C: Median speed linear mixed-effects modeling

To identify the impact of substrate on median walking speed, we used a linear mixed effects model (lmer function, lme4 package) [1]. The normality of the median speed data was tested using multiple transformations, with the best result using the square root of the speeds. An independent factor model:

sqrt(v_med) ~ substrate + day + (1+substrate|colony)

was compared to a null model:

sqrt(v_med) ~ day + (1+substrate|colony)

and a model that included interactions between the substrate and colony categorical factors:

sqrt(v_med) ~ substrate * day + (1+substrate|colony)

The assumptions of the linear-mixed effects model were tested by checking the normality of the model residuals and the linearity of the normal Q-Q plot of the residuals.

Chi-squared Likelihood Ratio Tests comparing the independent-factor and null models determined whether substrate type significantly influenced median speed while accounting for variation due to the colony (i.e. how many ants were collected, caste breakdown of ants collected, slight weather or seasonal fluctuations) and the day of the recording trial (ants generally were slower during the recording session on the second day). Comparison of the independent-factor and interacting-factor models determined if the day of recording differentially influenced ant walking speed based on the substrate (Table S1). Comparisons between substrates were calculated by finding the estimated marginal means of the linear mixed effects model (emmeans package, tukey multiplicity adjustment).

Appendix D: Outdoor preference experimental design

The design of this set-up went through many iterations before accurately testing ant preference. Argentine ants lay pheromone trails to direct their nestmates to food sources. We found that when separating the flat and rough substrates (i.e. a Y- or T-preference test), stochasticity in early substrate exploration would ultimately determine ant preference. For example, if the first few ants to explore the setup found the food by walking over the rough substrate first, their pheromone trail would bias the rest of the recruited workers. In these circumstances, only a few ants would actually test preference, leaving a high noise:signal ratio. Instead, by placing the flat and rough rectangles directly next to each other, all ants could dynamically shift between flat and rough pathways while recruiting to the same food source.

Every morning between 7 and 10 AM, clean 3D-printed substrates and fresh food was placed in the set-ups and the paper covering for the plank was replaced. We randomly determined the ordering of the 1, 3, and 5 mm substrates, as well as their left or right position with respect to the paired flat substrate. Because ants demonstrate a strong navigational memory (54), we rotated each set-up by 180° each day. Each set-up was covered by a darkened plastic box with integrated LED lighting to normalize lighting conditions and prevent most other animals from discovering the food.

Appendix E: Processing time lapse videos of outdoor preference experiments

For every cantilever-bridge set-up, the location of each smooth and rough rectangle was manually defined on a video still frame. Any bright spots (pixel value < 100, i.e. reflections of LEDs) or dark spots (pixel value < 150, i.e. dirt on rectangle) that remained throughout 70% of the video were adjusted by adding or subtracting 50 pixels, respectively. For each frame, a background was calculated by averaging the pixel values of the two frames before and after that frame. A normalized video was produced by dividing each frame by its background image. Each normalized frame was thresholded to find relatively dark pixels (value > 1.15) and analyzed to find connected foreground objects (ndimage.find_objects). All objects larger than 6 pixels and smaller than 100 pixels were identified as ants. The number of identified ant pixels in each rectangle was summed and compared between smooth and rough substrates. Occasionally, the sun would shine through an opening in the set-up box, dramatically changing the lighting of the video and disrupting automated analysis. In these sunspot frames, many relatively dark pixels were identified. Ignoring any frames with more than 1500 ant pixels ensured that only sunspot frames were removed from the analysis. The total percentage of ant pixels on smooth vs. rough substrates was calculated and compiled for each day of recording.

Appendix F: Deep-learning tracking of ant landmarks

To implement LEAP tracking on footage of highly-variable ant walking, we (1) produced trustworthy input ant-centered videos, (2) hand-digitized an extensive training set of frames, and (3) implemented post-tracking confidence checks to improve or remove untrustworthy data.

1) Generating ant-centered videos

During the full-body analysis, centroids were fit to each identified ant, identifying the angular orientation of the body but without specifying the directionality of the head. To generate videos cropped around the ant, first required a robust method for finding the head of the ant in each frame (Fig S6A). All processing was conducted using custom scripts in Python.

A first estimate of the head direction of each ant was determined by attempting to identify the antennae. For each ant-sized contour of interest in a thresholded frame (generated in the full-body automated tracking), this process involved: (a) removing all other contours from the image, (b) removing all legs and antennae (distance transform < 4) to fit a rectangle around the body points, (c) for each leg/antennae pixel, finding the distance from the rectangle center along the long-axis of the rectangle (ant body), (d) comparing the sum of the parallel distances to all the leg/antennae pixel on each side of the rectangle (the two ends of the ant). In situations where there was greater than a 10% difference in the sum, the end with a greater value was identified as the head. When there was less than a 10% difference, the original contour fitting angle was used as the ant facing angle. This method assumes that the anterior placement of the antennae results in more appendage pixels in front of vs. behind the ant. While this is often true, variation in the positions of the antennae and hindlimbs can obscure this method, requiring further analyses to confidently identify the facing.

Once ant-head-identification had been improved for each frame, facing angles were compared across each trackway to remove any aberrant angles. Unusual body positioning of the ant caused some incorrect identifications (mistaking the ant gaster as the head) or, if the ant-head-identification did not result in an improved estimate, the original contour fitting method did not take into account the directionality of the ant's facing. To ensure that the facing of the ant was consistent, the ant's facing angle was compared across all frames in each trackway longer than 50 frames. Sections of consistent tracking of ant facing were defined by when the change in the sine or cosine of the ant's facing angle varied by less than 1. If the sine or cosine jumped by more than 1 between frames, this was due to a 180° rotation with the other side of the ant (head or gaster) now being used to identify the facing. Every other section was then corrected by inverting the sine and cosine traces. These traces were then smoothed using a moving average (custom script, window size = 11 frames, ignoring any windows with fewer than 2 non-nan values), then used to calculate the new ant facing angle in every frame. Using this method ensured that the same side of the ant was tracked throughout the whole trackway.

Each trackway was now used to generate ant-centered images with a dark background. Each background-divided frame was rotated so that the direction of the ant facing aligned with the +x direction (imutils.rotate_bound, Python) and cropped to a shape of 200x200 pixels around the center of the ant. Each cropped, background-divided image was then inverted and adjusted to increase the contrast between ant and background. Having undergone background-division, the pixel values of each image ranged from 0 to a positive value less than 255 (often closer to 20-30) with 1 corresponding to pixels with the same value as the background and values <1 corresponding to pixels relatively darker than the background, including the ant. Each image was inverted (pixels previously darker than the background are now brighter, >1) and shifted so that the now relatively darker pixels range from 0 to 5 and the now relatively brighter pixels (including the ant) range from 5 to 255 pixels.

The LEAP tracking program will identify the most likely pixel that resembles each point tracked in the training set, meaning that the presence of other ants within an ant-centered frame will significantly disrupt tracking. To remove these other ants, each ant-centered frame was thresholded (cutoff = 25), morphologically closed (OpenCV, kernel = cross, iterations = 1) to find all contours. Small contours (<10 pixels) and any contours touching the edge of the image were set to a pixel value of 3. If the central centroid was removed in this process (for example, if the ant of interest was touching another ant) or if the center of the ant was within 50 pixels from the edge of the full, non-cropped frame, the image was set to black.

At this point, we had generated a series of ant-centered images with the ant facing the same direction for all frames. However, some trackways were either too short or included ant positions particularly difficult for reliable head identification, resulting in a trackway that followed the tail and not the head of the ant. A support vector machine (SVM) was used to identify these completely flipped trackways. A training group of 1285 ant-centered pictures were used to train the SVM to classify 3 ant facing categories: right (ant faces in +x direction), left (ant faces in -x direction), or blank (the ant-centered picture is completely black as would be generated if the ant facing angle = nan). A principal components analysis applied to the training set generated a set of 15 eigenimages, which were then used to compress the input image data. Using a practice test set of 25% of the training set, the SVM (sklearn package, C = 1000, gamma = 0.0001, kernel - ‘rbg’, class_weight = ‘balanced’) identified left and right ant images with 99.6% accuracy. The first 100 frames of each trackway were fed into the SVM and classified as right, left, or blank. If the number of left-facing images outnumbered the right-facing images, the ant facing angles and ant-centered images for the trackway were rotated by 180°. The images for each trackway were saves as a .h5 file to be compatible with the LEAP analysis functions.

2) Training the LEAP tracker

The LEAP workflow uses a training set of tagged subject-centered images to predict point placement in novel images. Our skeleton of points-to-track included 10 points: the distal tarsus of each limb, 2 points along the mid-sagittal axis of the thorax, and the tip of each antenna. In preliminary tests we were able to track 36 points (including 4 points on each limb), but we ultimately limited the tracked points to reduce tracking time and file size.

We generated a network training set by iteratively tracking a large dataset of non-stereotyped walking movements (Fig S6B). We could reliably track walking on flat ground using a training set of only a few dozen flat substrate images, but to accurately track the variable movements on uneven substrates required a much larger training set. Our final model used 679 frames (the maximum number before causing memory errors with a NVIDIA Geforce GTX 1080 8GB GPU), specifically focusing on tracking unusual limb movements observed on 1 and 3 mm substrates. The LEAP network was trained using the following settings: scale = 1, kernel for confidence maps = 5, mirrored images enabled, leap_cnn network architecture, 64 base filters, 5 deg rotation angle, 25 epochs, 50 batches per epoch, 50 samples per batch, validation fraction = 0.1, AMSGRAD enabled, learning rate reduction factor = 0.1 after 3 epochs with the learning rate changing by less than 1e-5. The final network training took approximately 30 minutes, with a learning rate loss of 4.35e-5 after 25 epochs.

Each trackway's .h5 file generated in section (2) was run through the prediction model, requiring between 3 and 30 seconds depending on the number of frames.

3) Post-processing and validation of LEAP tracked data

Despite our efforts to generate trustworthy input ant-centered videos and a robustly-trained deep learning network, occlusions of the limbs and general kinematic variability resulted in some inaccurate tracking predictions. Identifying and removing these points was a multistep process.

First, untrustworthy tracked data points were identified and removed. In addition to predicting the tracked points in each image, the LEAP program also provides a confidence value, ranging from 0 to 1, associated with each tracked point. After visually investigating how the confidence value corresponded to tracking accuracy, we chose a confidence cutoff of 0.6, removing any tracked values with a confidence below this value (Fig S7). To account for other tracking inaccuracies, we applied two additional conditions. (1) For the trace of each tracked point (with respect to the ant), any sections of tracking that were bound by jumps of over 10 pixels were removed. (2) Any outlier points were removed. We identified outliers by first determining the "middle range" of the x- (anterior-posterior) and y- (medial-lateral) coordinates for each trace using the middle 50% of values in each axis. If the middle range was less than 90% of the full range, outliers were calculated by finding any point that differed from the x or y mean of the middle range points by more than ⅔ of the middle range.

Second, each tracked trace was filtered to remove noise. After interpolating any small gaps (<= 5 nan values in a row), each tracked section of a trace longer than or equal to 9 values was low-pass filtered (Scipy, butterworth, n=2, $\omega_n$ = 0.3). Sections of fewer than 9 values were set to nan. If after post-processing, fewer than 50 finite (non nan) data points remained in the full trace, that trace was removed from further analysis.

Appendix G: Identifying tarsal touchdowns

Because feet stay mostly in place during stance (with respect to the lab coordinate system), foot speed can be used to identify tarsal stance location. First, periods when tarsal speed was below a speed cutoff were identified (Fig S8). The speed cutoff was either 1, 1.5, 2, or 3 pix/fr based on the extent of tracking noise for the joint (the median speed of the time series). Second, any periods when the foot was below the speed cutoff for more than 5 frames were considered a stance (a gap of 1 or 2 non-slow or nan points was allowed). Stance locations were found by averaging the x- and y-locations of the foot during the first 5 frames of stance. Third, any stance locations within 10 pixels of the previous stance were removed. Lastly, TD timing was determined by searching within the 12 frames before an identified stance to find when the foot first approaches within a set number of pixels from the stance location. The pixel cutoff was either 2 or 3 pixels based, again, on the extent of tracking noise for the joint. Any TDs without at least 3 tracked frames before the TD frame were not trusted and removed from further analysis.

Two variables were calculated for each trusted TD.
(1) The location of the foot at the time of TD was related to the full frame based on the original estimates of ant location and orientation (centroid and heading), which were used to generate the ant-centered images input to LEAP.
(2) The location of the foot relative to the ant at the time of TD was calculated using the LEAP-tracked location of the ant thorax, defining the mid-sagittal axis using the relation of the tracked neck point to the thorax.

Strides were identified as occurring between two trusted TDs and with the following two conditions. First, 80% of the stride had to be tracked to ensure that no intermediate TDs were missed. Second, other than the continuous periods of slow foot speed at the beginning and end of a stride, only one datapoint could dip below the slow speed cutoff (Fig. S8). We validated these metrics by plotting TD timing on 17 videos (including 14 videos on non-flat substrates) and visually checking for accuracy.

Appendix H: Average stride speed linear mixed-effects modeling

We selected input data for the models from stereotypical straight strides on flat ground. To identify stereotypical strides, we applied a kernel density estimate (sklearn.neighbors.KernelDensity, bandwidth using Scott's Factor) to the speed versus stride frequency graph, exporting only strides with a density > 0.005. In all, 21152 straight walking strides were used to generate the linear models.

After checking the normality of the average velocity variable, three models were generated from the data. All three models were a random-intercept, linear mixed effects model with the following equations (lme4::lmer function in R)[1]:

1. The "Full" model
   lmer(ave_v ~  stride_freq + stride_len + day + (1|colony)
2. The "constant stride length" model
   lmer(ave_v ~  stride_freq + day + (1|colony)
3. The "constant stride frequency" model
   lmer(ave_v ~  stride_len + day + (1|colony)

The goodness of fit of these models was confirmed by examining quantile-quantile and residual plots (**Fig. S*****). Although the quantile-quantile plots for the full and constant stride frequency models displayed light tails, this derives from our method of selecting stereotypical strides.

Some strides were well predicted by these models, while other diverged largely. To isolate just the well-modeled "inlier" strides, we applied a kernel density estimate to the full model error distributions (R: density function, kernel = gaussian). All strides with a density greater than 0.01 were used to calculate a mean and standard deviation on each substrate. Inlier strides were identified as those with full model prediction errors within 4 standard deviations of the mean. Outlier strides had errors greater than 4 standard deviations. These calculations were only performed for the full model, but the inlier and outlier stride classifications were applied to all models.

**Figure S1.**

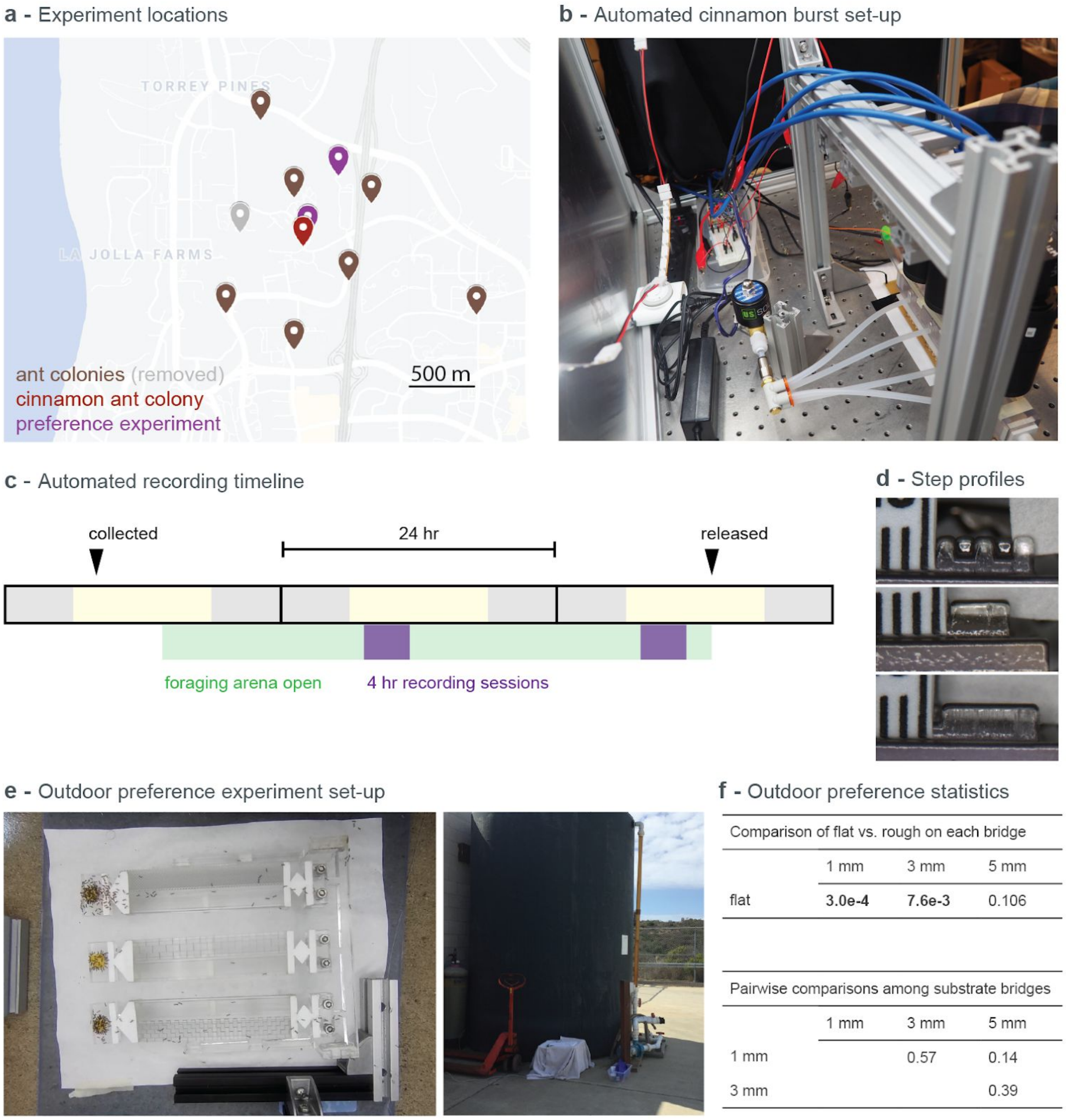


Comparison of flat vs. rough on each bridge

| | 1 mm | 3 mm | 5 mm |
|---|---|---|---|
| flat | **3.0e-4** | **7.6e-3** | 0.106 |

Pairwise comparisons among substrate bridges

| | 1 mm | 3 mm | 5 mm |
|---|---|---|---|
| 1 mm | | 0.57 | 0.14 |
| 3 mm | | | 0.39 |

**Figure S1.** Experimental procedures and outdoor preference statistics. (a) Locations of collected ant colonies and outdoor preference experiment in La Jolla, CA. (b) An arduino-controlled solenoid valve released cinnamon-infused air into the ant tunnel above each substrate. (c) Experimental timeline for normal and cinnamon recording trials. (d) Magnified views of each 3D-printed substrate used to analyze the roundedness of the checkerboard edges. (e) Photos were recorded every 3 min for 24 hours of a custom cantilever support structure that forced foraging ants to walk over a flat or rough substrate. (f) Ant significantly prefered flat versus rough substrates for 1 and 3 mm substrates (bold, $p < 0.01$) but comparisons among substrates showed no significance.

**Figure S2.**

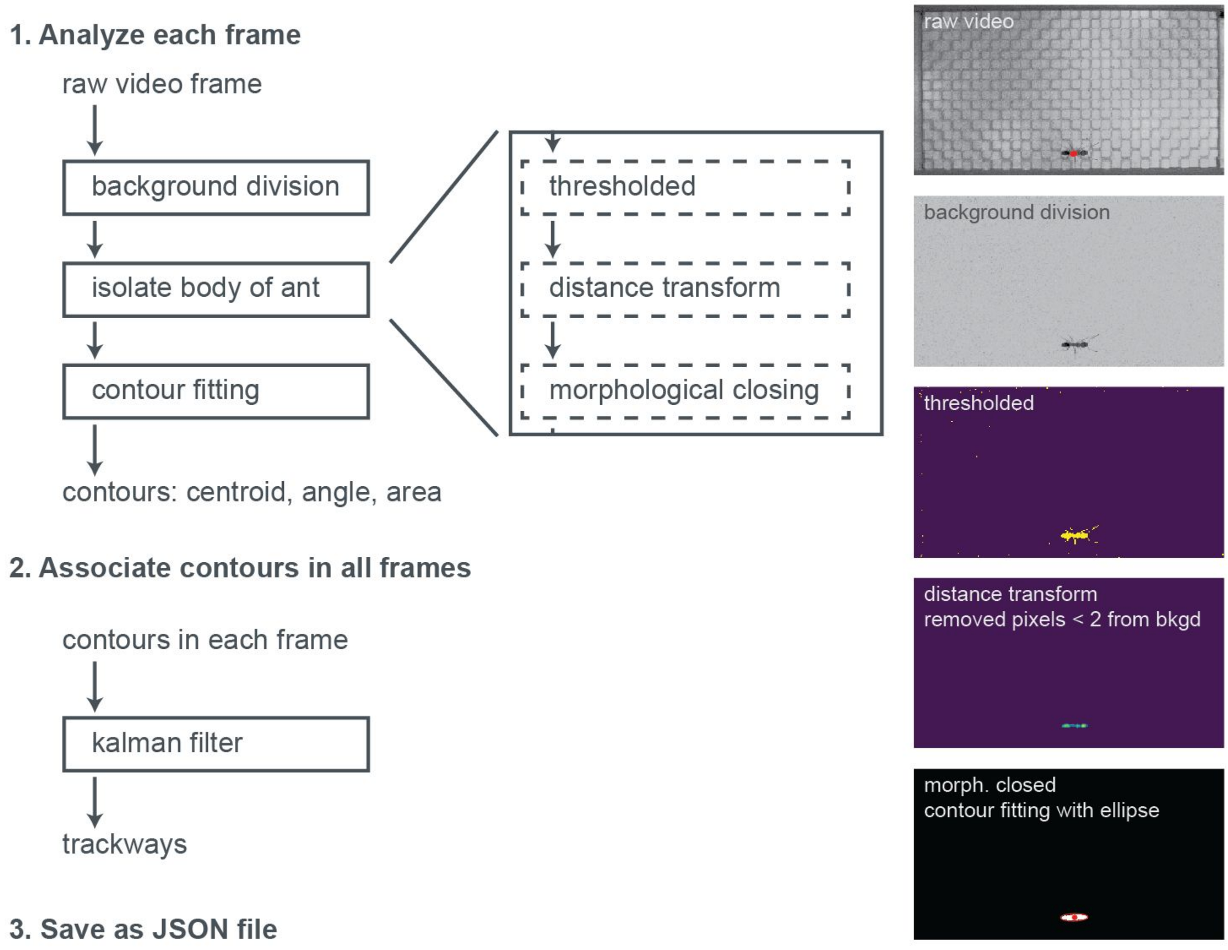


**Figure S2.** Analysis workflow of full ant body tracking, implemented in Python. Every frame of each video was analyzed individually to identify the location and approximate orientation of each ant. Then a kalman filter identified ants conserved across subsequent frames, producing ant trackways.

**Figure S3.**

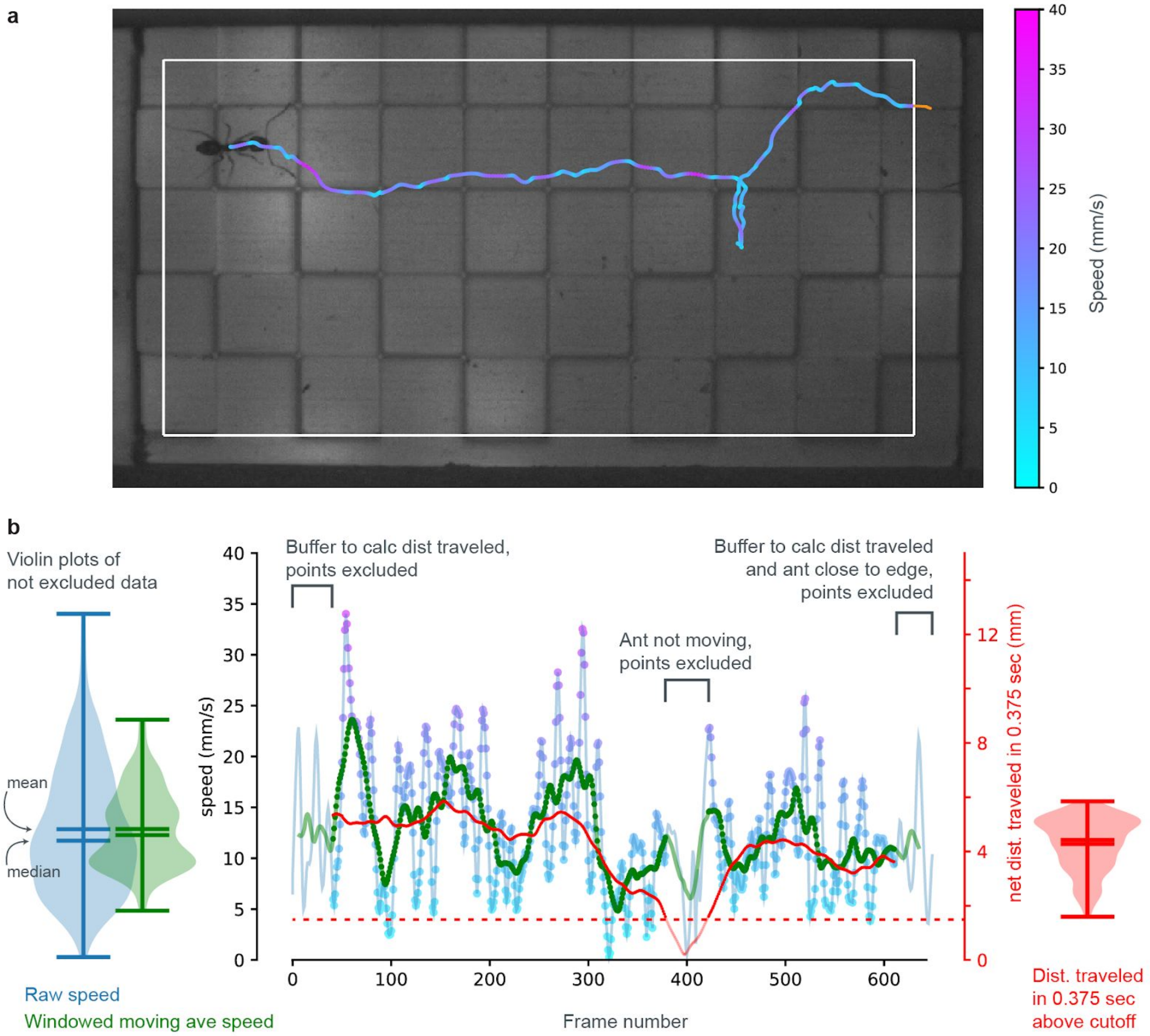


**Figure S3.** Processing of instantaneous walking speed data. (a) Any frames where the center of the ant is within 60 pixels (~1.9 mm) of the edge of the frame (white line) are removed from analysis (orange trace). (b) The net distance traveled in 90 frames (0.375 s) (red) was used to remove frames when the ant is stopped. A cut-off of 50 pixels (1.56 mm) identified times of stillness. The raw speed trace (blue to purple points) for trusted points was used to calculate the trailway median speed (blue violin plot). A windowed moving average (window = 0.1 sec) was also calculated (green trace and violin plot).

**Figure S4.**

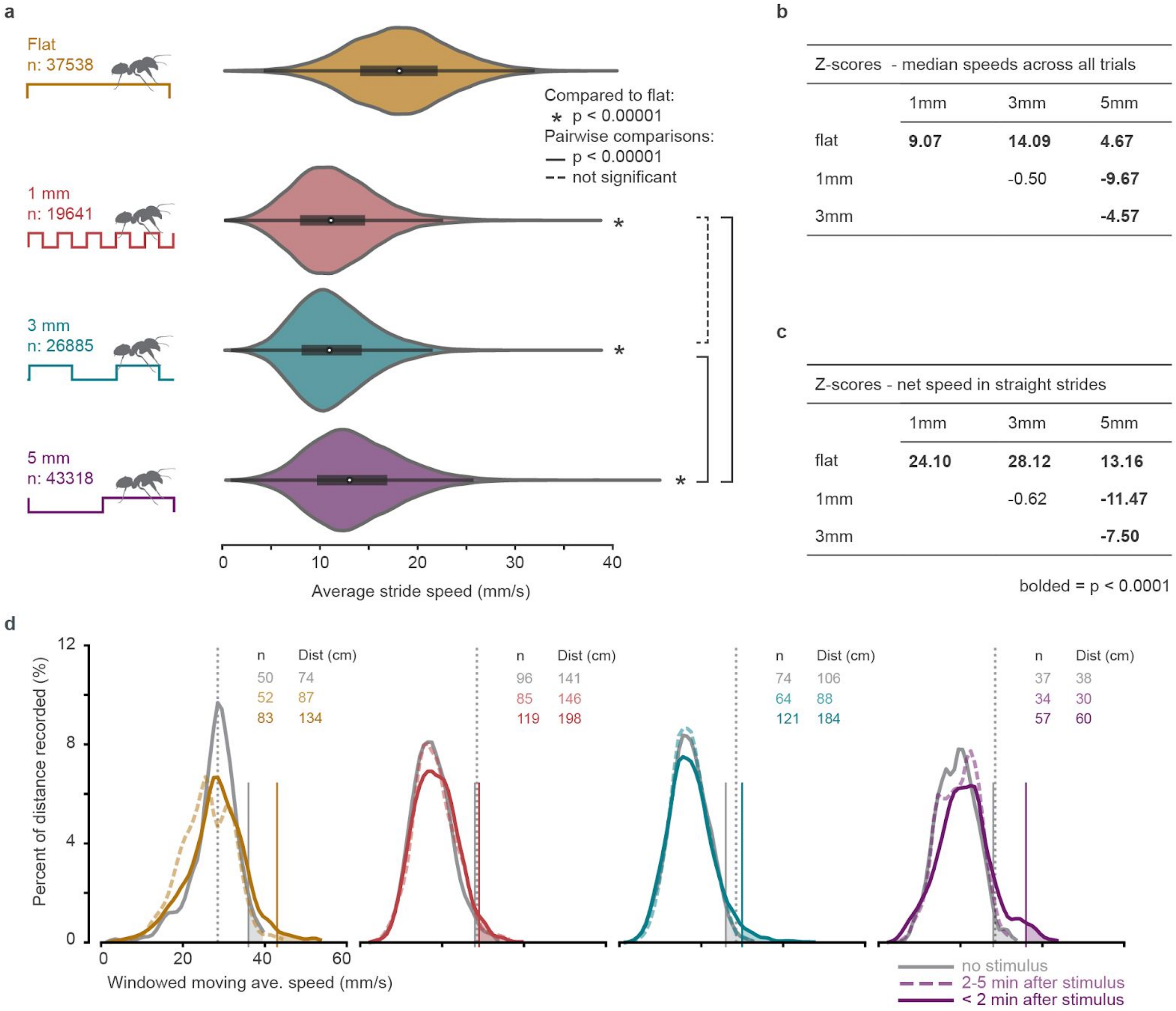


**Figure S4.** Ant walking speeds during straight walking and during noxious stimulus trials. (a) For all walking strides with the ant moving straight (heading < 15°) the net body speed distribution was approximated by a gaussian kernel density estimate. Statistically significant differences among substrates (bold) for the analysis of median walking velocity across all trials (b) mirrored findings using only straight strides (c). (d) The distance traveled at each speed in one colony without any intervention (solid gray lines), within 2 minutes of an air burst (solid, colorful lines), and between 2 and 5 minutes after a burst (dashed, colorful lines). Vertical lines show the flat ground, unperturbed median speed (gray dashed) and peak speed cutoff at 98% of the cumulative distance traveled for unperturbed (gray solid) and noxious stimulus (solid colorful) conditions.

**Figure S5.**

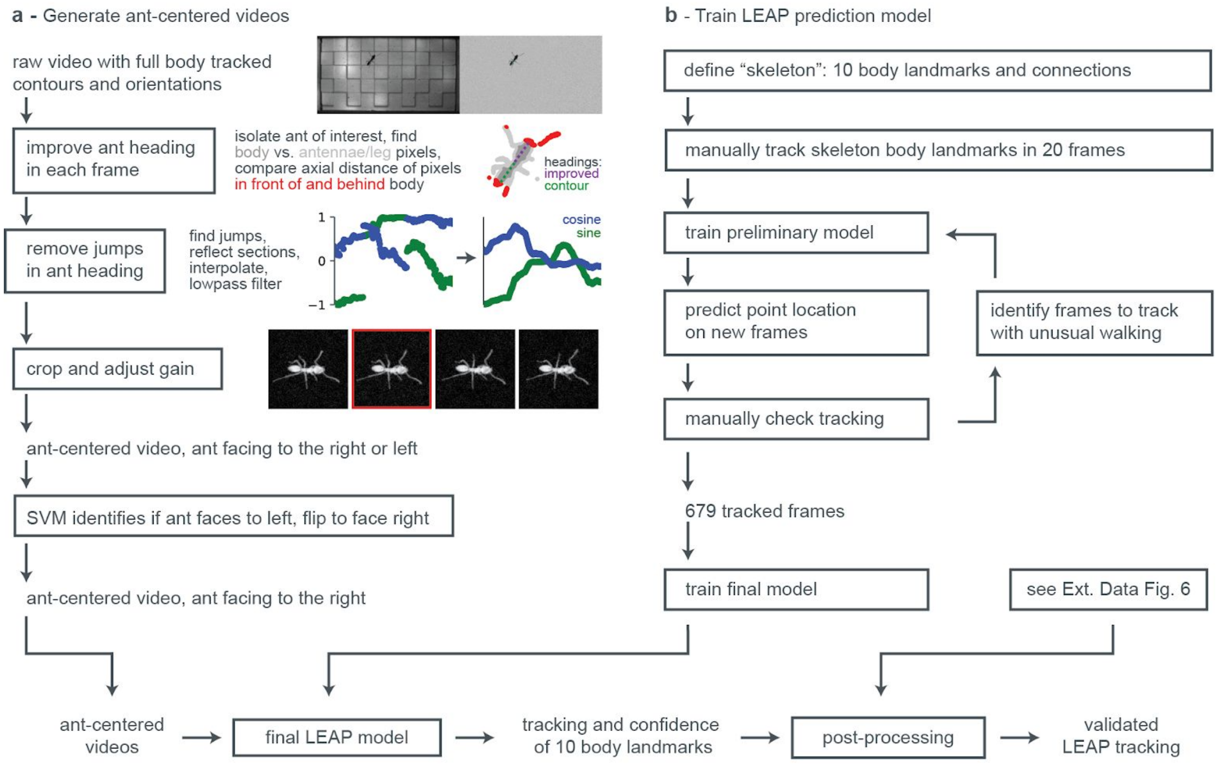


**Figure S5.** Workflow for deep learning (“LEAP”) tracking of limbs, body, and antennae. First, ant-centered videos are generated by reliably identifying the facing of the ant in each frame, then cropping and rotating each frame around the ant, adjusting the gain of the background and foreground to enhance contrast, and lastly using a support vector machine (SVM) to ensure that the ant faces to the right. The LEAP prediction model is developed through iterations of manually tracking points, training a model, and predicting tracked locations on new frames. The LEAP prediction model outputs tracked points in each ant-centered video frame.

**Figure S6.**

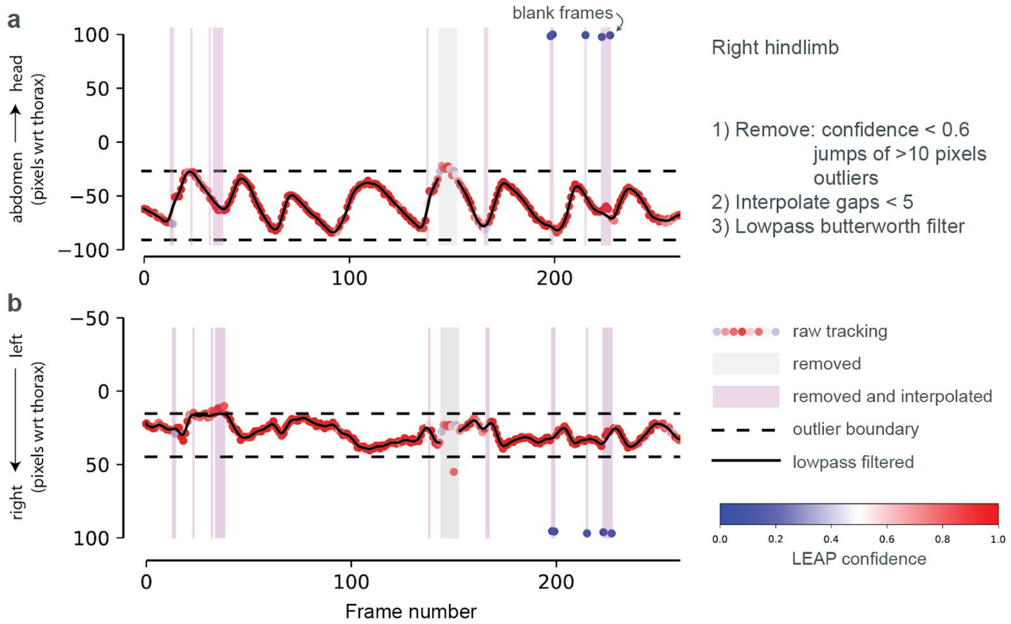


**Figure S6.** Post-processing of raw LEAP tracking for the right hindlimb tarsus in the 3mm trackway depicted in External Data Fig. 3. Untrustworthy tracking is identified with 3 conditions based on the (a) anteroposterior and (b) mediolateral traces: a LEAP confidence value <0.6 (blue coloration), a jump in location of more than 10 pixels, a location outside a cutoff range determined using the middle 50% of locations in the trial (dashed lines). After untrusted data are removed, any gaps < 5 values are interpolated, and sections of tracking > 9 values are lowpass filtered.

**Figure S7.**

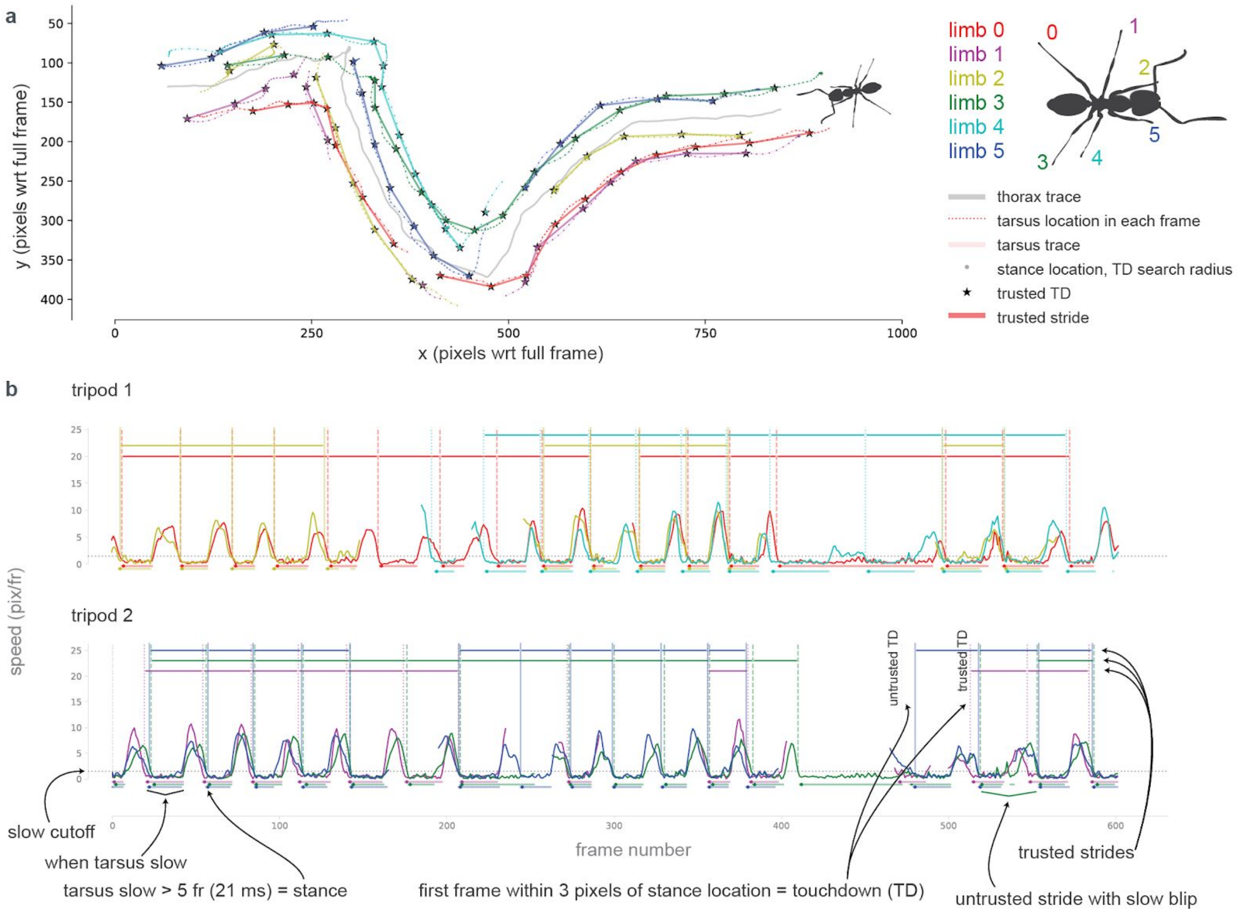


**Figure S7.** Identification of touchdowns (TDs) and strides using tracked tarsus data. (a) Tarsus location relative to the full frame shows untrusted (circle) and trusted (star) locations. (b) Sections of total instantaneous tarsal speed with 5 or more consecutive datapoints below a cutoff (gray dashed line) indicate a stance (circle). Stance location is calculated from the first five frames, and used to find TD timing. Trusted TDs have at least 3 frames tracked before the TD. Trusted strides connect consecutive pairs of trusted TDs with more than 80% of the stride tracked and without a blip of slow frames within the stride.

**Figure S8.**

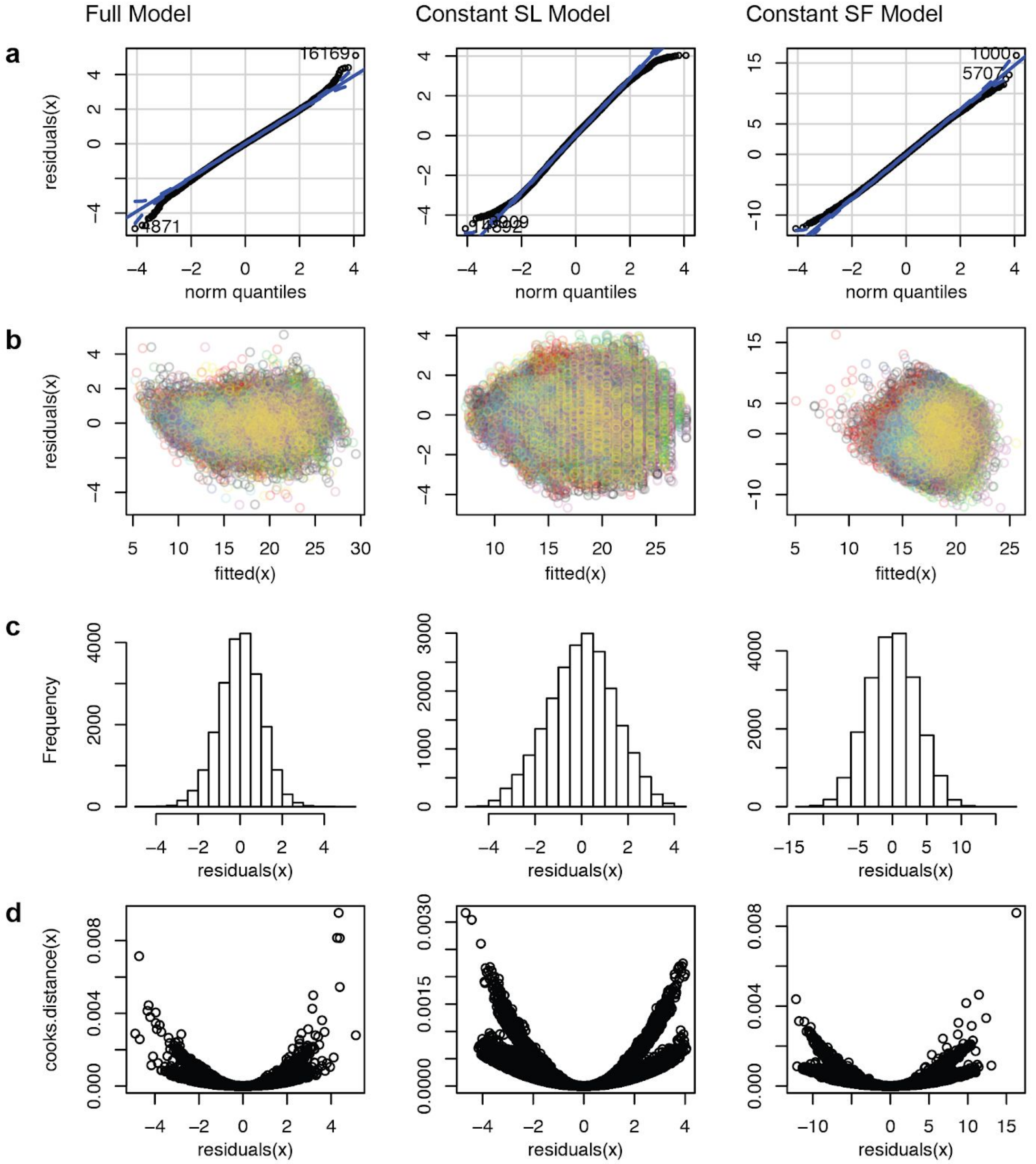


**Figure S8.** Fit of the full, constant stride length (SL), and constant stride frequency (SF) models generated from flat walking strides. (a) Quartile-quartile plots show a primarily linear relationship with light tails. These tail abnormalities are anticipated given our method of selecting "stereotypical" flat walking strides. (b) Plots of residuals versus fitted values show limited structure. Colors represent different colonies. (c) Histograms of the residuals show an approximately normal distribution. (d) Cook's distance versus residual plots show an approximately symmetrical influence of large magnitude residual points.

**Figure S9.**

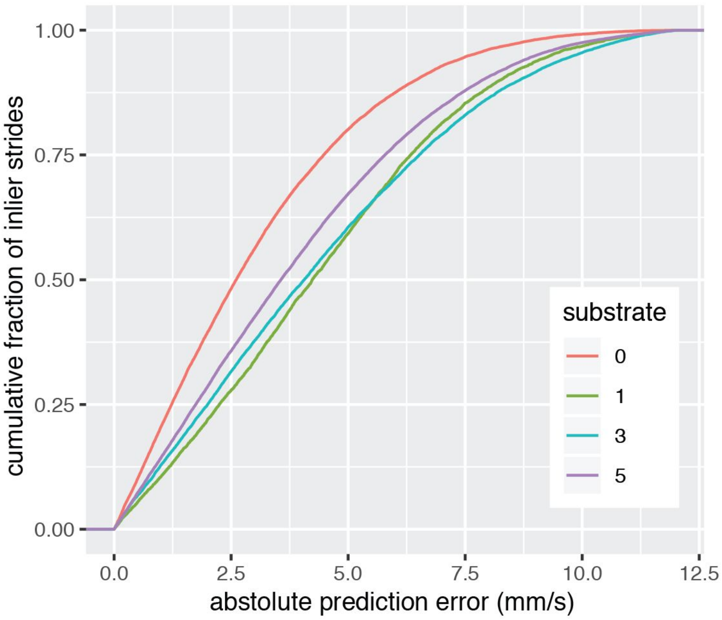


**Figure S9.** Cumulative distribution of inlier strides predicted by the constant stride frequency model. The y-axis shows the fraction of all inlier strides within the prediction cutoff along the x-axis. Each line represents a different substrate.

**Figure S10.**

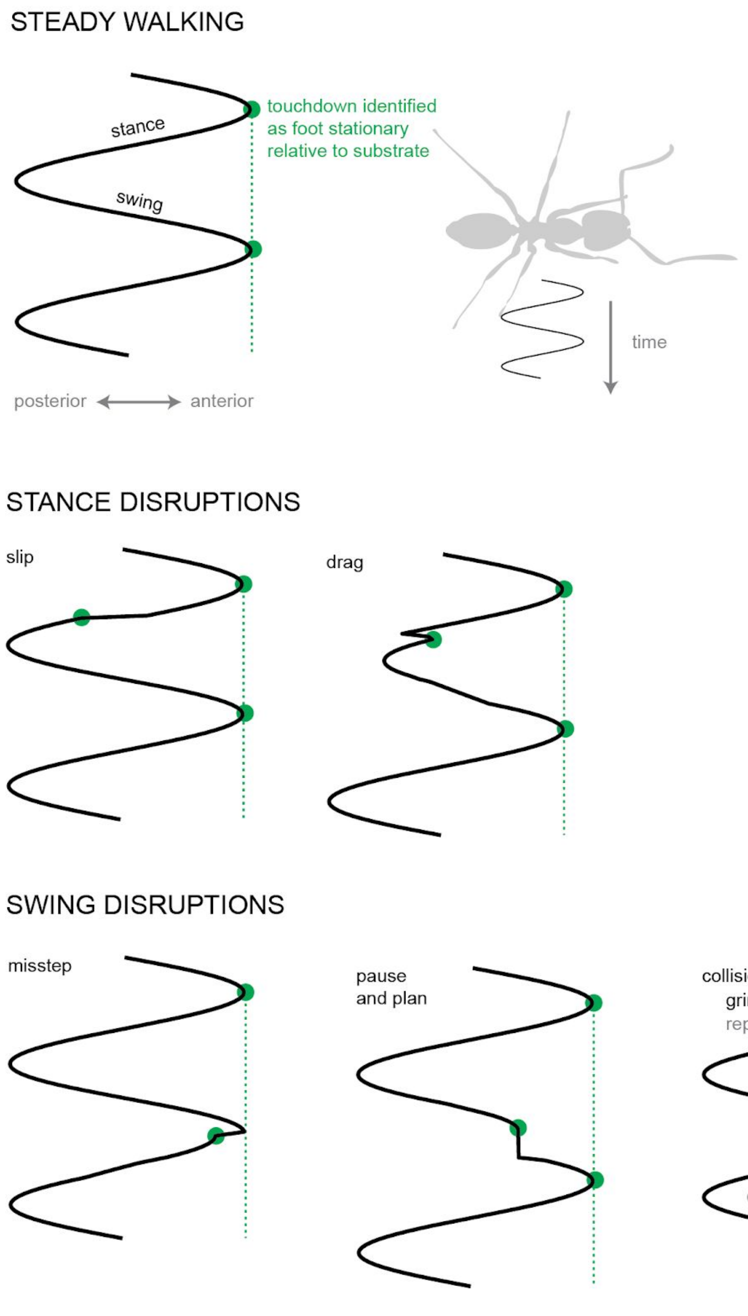


**Figure S10.** Diagrams of outlier stride classification types. The foot trajectory relative to the ant oscillates anteriorly and posteriorly (x-axis) across time (y-axis). Green dots represent times when the foot touches the ground to begin stance, during which the foot remains in one place relative to the global coordinate system. Disruptions of steady walking during stance include foot slips and drags. Disruptions of walking during swing include scenarios when foot touchdown is temporally delayed (missteps) or early (“pause and plan” or collisions). “Pause and plan” strides result from the ant temporarily tapping its limb along the ground before finding a final placement, often occurring at obstacle edges.

**Table S1.** Description of linear-mixed effect model parameters for full, constant stride length (SL), and constant stride frequency (SF) models. For each model, t-values and p-values are listed showing the relative influence of each factor, though a more rigorous chi-squared likelihood ratio test compares each reduced model with the full model. The full model also includes correlation values, confirming that stride length and stride frequency are not tightly correlated.

| Fixed effects: | intercept | SF | SL | day |
|---|---|---|---|---|
| **Full model** | | | | |
| t-value | -116.6 | 491.9 | 145.3 | -6.5 |
| p-value | < 2e-16 | < 2e-16 | < 2e-16 | 6.47E-11 |
| corr. with SF | -0.304 | | -0.202 | 0.097 |
| corr. with SL | -0.645 | -0.202 | | 0.052 |
| **Constant SF model** | | | | |
| t-value | 5.0 | | 70.7 | -15..5 |
| p-value | 0.000882 | | < 2e-16 | < 2e-16 |
| **Constant SL model** | | | | |
| t-value | -16.1 | 376.2 | | -10.0 |
| p-value | 4.51E-08 | < 2e-16 | | < 2e-16 |
| **Likelihood Ratio Chi-Square test of fixed effects** | | | | |
| chi value | | 53301 | 14644 | 42.718 |
| p-value | | < 2.2e-16 | < 2.2e-16 | 6.32E-11 |

**Legends for Movies S1 to S2**

Supplementary Video 1

Wild ants walking over smooth or rough substrates to reach a food source. (Left) Raw footage of the outdoor preference set-up with ants identified in red (1 mm bridge), green (3 mm bridge), and blue (5 mm bridge). (Center) Isolated regions of the bridges were background subtracted using the neighboring frames. Non-touching blobs were identified as ants. (Right) The number of ant-pixels identified on the flat or rough substrate for each bridge.

Supplementary Video 2

Post-processing of limb tracking for an ant walking on a 3 mm substrate. (Left) Limb, antennal, and body landmarks generated from the deep-learning based tracker are projected onto a background-subtracted cropped view. A large white dot represents the approximate ant body location from the "Automated full-body tracking" workflow. The smaller dots represent LEAP tracked landmarks, with color representing the output confidence. (Center) Post-processed tracking of body (white), left tarsi (blue), and right tarsi (magenta) points. White circles represents a trusted touchdown. Trusted strides have a white line connecting the tarsus with its prior touchdown location. (Right above) The full recorded frame. (Right below) Time-varying total velocity of each tarsus. Dots represent trusted touchdowns.
